# Transcription-specific synaptic plasticity mediates engram computations

**DOI:** 10.64898/2026.08.22.746450

**Authors:** Douglas F. Tomé, Meizhen Meng, Xiaochen Sun, Li Yao, Tim P. Vogels, Yingxi Lin, Claudia Clopath

## Abstract

Memory is encoded by sparse ensembles of neurons. While activity-dependent transcription and learning-induced synaptic plasticity are causally related to memory formation and recall, the link between the transcriptional regulation of synaptic plasticity and memory computations remains elusive. In particular, even though transcriptionally defined neuronal ensembles within a memory engram exhibit specific forms of synaptic plasticity and support distinct behavioral outputs, it is still unclear whether transcription-specific synaptic plasticity drives ensemble computations, rather than merely serving as a marker of ensemble identity. Here, we demonstrate that transcription-specific synaptic plasticity enables ensemble computations essential for learning-induced adaptive behaviors. In the mouse dentate gyrus (DG), we found that neuronal ensembles genetically defined by *Fos*-dependent transcription engage plasticity in feedforward excitatory synapses, whereas those defined by *Npas4* -dependent transcription engage plasticity in recurrent inhibitory synapses. We modeled spiking neural networks with transcription-specific synaptic plasticity and observed that *Fos*- and *Npas4* -dependent ensembles emerged following learning and stabilized with memory consolidation. Our computational model predicted that blocking *Fos*- or *Npas4* -dependent synaptic plasticity disrupts memory generalization and discrimination, respectively. By acutely deleting *Fos* or *Npas4* in the DG to selectively block *Fos*- or *Npas4* -dependent synaptic plasticity, we conducted contextual fear conditioning experiments whose results supported our computational model’s prediction. Our study provides causal evidence that specific transcriptional programs induced in distinct neuronal ensembles differentially engage synaptic plasticity and thereby regulate memory computations.

## Introduction

An engram refers to the enduring physical changes in the brain that preserve the memory of a specific experience.^1–3^ For instance, sparse neuronal ensembles activated during memory acquisition remain necessary and sufficient for recall,^4, 5^ and learning-induced synaptic plasticity is essential for memory formation.^6–8^ For synaptic changes to persist over the timescales of long-term memory, neuronal activity must engage the cell nucleus to induce new gene transcription.^9–11^ This activity-dependent transcription regulates synaptic plasticity through the induction of genes encoding effector proteins that build, remodel, and tune synapses (e.g., *Arc* and *Homer1a* regulate the strength of excitatory synapses^12–14^). Loss- and gain-of-function manipulations of various activity-induced genes regulating synaptic plasticity (e.g., *Arc, Bdnf, Creb*, and *Mef2*) have been shown to disrupt memory formation and recall in behavioral paradigms such as associative and spatial learning.^15, 16^ Together, these findings have linked the transcriptional regulation of synaptic plasticity to memory encoding and storage.

Our previous work found that a single memory engram encompasses transcriptionally defined ensembles that display specific forms of synaptic plasticity and are functionally distinct.^17^ Specifically, a contextual fear memory engram in the mouse dentate gyrus (DG) contains ensembles genetically defined by the transcriptional pathways downstream of the immediate-early genes (IEGs) *Fos* and *Npas4*. The *Fos*-dependent ensemble exhibits excitatory synaptic plasticity and promotes memory generalization, while the *Npas4* -dependent ensemble exhibits inhibitory synaptic plasticity and promotes memory discrimination. In effect, rather than a single transcriptional program acting uniformly within an engram, distinct programs partition the engram into ensembles with specific synaptic and computational properties. Notably, generalization and discrimination are opposite and complementary ensemble computations that must be balanced to enable both recognizing shared features across related experiences and distinguishing between different events.^18–22^ Critically, knowledge of the neurobiological mechanisms that mediate the generalization-discrimination balance may shed light on adaptive learning as well as maladaptive conditions in which this balance is compromised, such as post-traumatic stress disorder (PTSD) and panic disorder.^23, 24^ Yet, whereas prior work provided causal evidence linking synaptic plasticity to memory storage,^6–8^ the link between transcriptionally regulated synaptic plasticity and the generalization-discrimination balance has so far remained correlational. As such, it is still unclear whether transcription-specific synaptic plasticity actively drives ensemble computations, rather than merely marking ensemble identity.

To investigate the link between transcription-specific synaptic plasticity and neural computations, we first examined which excitatory and inhibitory synaptic inputs are selectively recruited by *Fos*- and *Npas4* -dependent ensembles in the DG following contextual fear conditioning (CFC). We found that *Fos*-dependent ensembles engage excitatory plasticity in feedforward synapses from the medial entorhinal cortex, while *Npas4* -dependent ensembles engage inhibitory plasticity in recurrent synapses from local cholecystokinin-expressing interneurons. Informed by our experimental results, we modeled spiking neural networks with transcription-specific synaptic plasticity by having half of the population of excitatory neurons engage plasticity only in feedforward excitatory synapses (*Fos* ^+^ neurons) while the other half engage plasticity only in recurrent inhibitory synapses (*Npas4* ^+^ neurons). Besides providing insights into the emergence and consolidation of *Fos*^+^ and *Npas4* ^+^ ensembles, our model predicted that blocking *Fos*- or *Npas4* -dependent synaptic plasticity disrupts generalization and discrimination, respectively, without impairing recall. Our experimental results supported our model’s prediction: acute deletion of *Fos* or *Npas4* in the DG blocked ensemble-specific excitatory and inhibitory synaptic plasticity, selectively impairing contextual fear memory generalization and discrimination, respectively. We therefore provide both computational and experimental evidence establishing that distinct neuronal ensembles within an engram recruit specific transcriptional programs that differentially engage synaptic plasticity to drive memory computations.

### Transcription-specific expression of synaptic plasticity within an engram

Our previous study showed that CFC recruits neuronal ensembles in mouse DG that are genetically defined by *Fos*- or *Npas4* -dependent transcription.^17^ The study also reported that *Fos*-dependent ensembles express excitatory but not inhibitory synaptic plasticity, while *Npas4* -dependent ensembles express inhibitory but not excitatory synaptic plasticity. Furthermore, the study examined which excitatory synaptic inputs onto *Fos*-dependent ensembles were selectively strengthened by electrically stimulating three major afferent pathways of the DG: the lateral perforant path from the lateral entorhinal cortex (LEC), the medial perforant path from the medial entorhinal cortex (MEC), and the mossy cell fibers from the dentate hilus. In addition, the study examined which inhibitory synaptic inputs onto *Npas4* -dependent ensembles were selectively strengthened by optogenetically stimulating gamma-aminobutyric acid (GABA)-ergic interneurons in the DG when pharmacologically inhibiting transmission from either parvalbumin-expressing (PV^+^) or cholecystokinin-expressing (CCK^+^) interneurons. Here, we sought to target specific excitatory and inhibitory synaptic inputs onto *Fos*- or *Npas4* -dependent ensembles using a combination of optogenetics and mouse genetics. Specifically, we trained mice in a CFC paradigm to induce the activation of either *Fos* or *Npas4* in sparse ensembles of DG granule cells. To identify *Fos*- and *Npas4* -dependent ensembles, we utilized our *Fos*-dependent Robust Activity Marking (*F*-RAM) and *Npas4* -dependent RAM (*N*-RAM) reporters.^17, 25^ For both *F*-RAM and *N*-RAM, the respective promoters drive the expression of a reporter gene (mKate2) under the temporal control of a modified doxycycline (Dox)-dependent Tet-off system.

We tested whether *Fos*- and *Npas4* -dependent ensembles express excitatory synaptic plasticity considering the three main excitatory inputs of the DG: LEC, MEC, and mossy cells in the DG hilus^26^ (Figure 1A-B). To selectively stimulate these excitatory inputs, we co-injected an AAV expressing a CaMKII-driven, light-activated channelrhodopsin (ChR2) into either the LEC or MEC, enabling optically evoked excitatory postsynaptic currents (oEPSCs) in DG granule cells by photoactivation of axonal terminals in the DG. We then performed simultaneous whole-cell patch-clamp recordings of oEPSCs from pairs of labeled and unlabeled neighboring neurons (*F*-RAM^+^/*F*-RAM^-^ or *N*-RAM^+^/*N*-RAM^-^) (Figure 1C). To stimulate excitatory inputs from the hilus, we injected an AAV expressing ChR2 unilaterally into the DG, labeling the hilar mossy cells whose axons project to the contralateral DG, where optogenetic stimulation and whole-cell patch-clamp recordings were carried out. We found that, compared to *F*-RAM^-^ neurons, *F*-RAM^+^ neurons exhibited significantly larger oEPSC amplitudes in response to stimulation of the MEC, while oEPSC amplitudes in response to stimulation of the LEC or the contralateral hilus remained comparable between the two groups (Figure 1D-E). Conversely, oEPSC amplitudes on *N*-RAM^+^ and *N*-RAM^-^ neurons were not significantly different (Figure 1D-E). Hence, *F*-RAM^+^ but not *N*-RAM^+^ neurons exhibit excitatory synaptic plasticity by strengthening their excitatory input selectively from the MEC.

**Figure 1.**
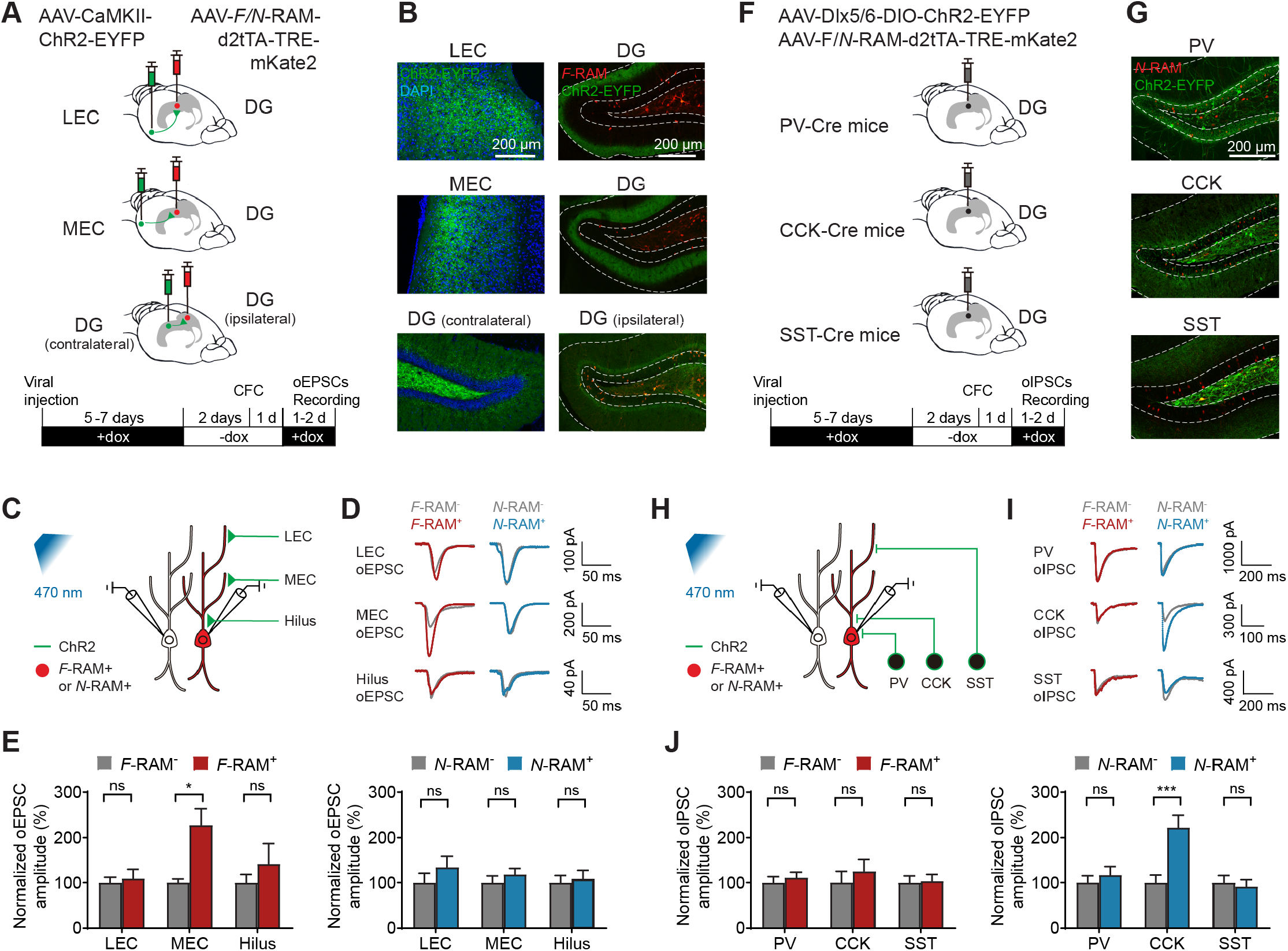
*Fos*- and *Npas4* -dependent ensembles differentially express synaptic plasticity. (A) Schematic of the experimental protocol to stimulate excitatory inputs. To target the LEC and MEC inputs, animals were injected with AAV-CaMKII-ChR2-EYFP to express ChR2 in the LEC or MEC, and also injected with AAV-*F* /*N*-RAM-d2tTA-TRE-mKate2 in the DG. To target the hilus inputs, AAV-CaMKII-ChR2-EYFP was injected unilaterally into the contralateral DG, which labeled the hilar mossy cells that send axonal projections to the ipsilateral DG. AAV-*F* /*N*-RAM-d2tTA-TRE-mKate2 was injected unilaterally into the ipsilateral DG, where whole-cell patch-clamp recordings were carried out (see Methods). (B) Representative images showing the expression of ChR2 and *F*-RAM^+^ neurons. (C) Schematic of the experimental protocol to simultaneously record oEPSCs on pairs of labeled and unlabeled neighboring neurons (*F*-RAM^+^/*F*-RAM^-^ or *N*-RAM^+^/*N*-RAM^-^) (see Methods). (D) Representative oEPSC traces from LEC (top), MEC (middle), and hilus (bottom) on pairs of labeled and unlabeled neighboring neurons. Left: *F*-RAM^+^/*F*-RAM^-^. Right: *N*-RAM^+^/*N*-RAM^-^. (E) Normalized oEPSC amplitude on pairs of labeled and unlabeled neighboring neurons. Left: *F*-RAM^+^/*F*-RAM^-^; *n* = 11 (LEC), 11 (MEC), 6 (hilus) pairs. Right: *N*-RAM^+^/*N*-RAM^-^; *n* = 8 (LEC), 8 (MEC), 8 (hilus) pairs. (F) Schematic of the experimental protocol to stimulate inhibitory inputs. To target specific inhibitory inputs, animals were co-injected with AAV-*F* /*N*-RAM-d2tTA-TRE-mKate2 and AAV-Dlx5/6-DIO-ChR2-EYFP into the DG of PV-, CCK-, or SST-Cre mice (see Methods). (G) Representative images showing the expression of ChR2 and *N*-RAM^+^ neurons. (H) Schematic of the experimental protocol to simultaneously record oIPSCs on pairs of labeled and unlabeled neighboring neurons (*F*-RAM^+^/*F*-RAM^-^ or *N*-RAM^+^/*N*-RAM^-^) (see Methods). (I) Representative oIPSC traces from PV^+^ (top), CCK^+^ (middle), and SST^+^ (bottom) interneurons on pairs of labeled and unlabeled neighboring neurons. Left: *F*-RAM^+^/*F*-RAM^-^. Right: *N*-RAM^+^/*N*-RAM^-^. (J) Normalized oIPSC amplitude on pairs of labeled and unlabeled neighboring neurons. Left: *F*-RAM^+^/*F*-RAM^-^; *n* = 9 (PV^+^), 8 (CCK^+^), 9 (SST^+^) pairs. Right: *N*-RAM^+^/*N*-RAM^-^; *n* = 13 (PV^+^), 16 (CCK^+^), 13 (SST^+^) pairs. Multiple Wilcoxon signed-rank tests with the two-stage step-up method of Benjamini, Krieger, and Yekutieli were performed on oEPSCs and oIPSCs. Data shown as mean ± SEM. \**q <* 0.05, \*\**q <* 0.01, \*\*\**q <* 0.001.

Next, we tested whether *Fos*- and *Npas4* -dependent ensembles express inhibitory synaptic plasticity considering the three main subtypes of GABAergic interneurons in the DG: PV^+^, CCK^+^, and somatostatin-expressing (SST^+^) cells.^27, 28^ To selectively stimulate these inhibitory inputs, we expressed ChR2 in different GABAergic subtypes by delivering AAV-Dlx5/6-DIO-ChR2-EYFP into the DG of PV-, CCK-, or SST-Cre mice (Figure 1F-G). We recorded optically evoked inhibitory postsynaptic currents (oIPSCs) from pairs of labeled and unlabeled neighboring neurons simultaneously (*F*-RAM^+^/*F*-RAM^-^ or *N*-RAM^+^/*N*-RAM^-^) (Figure 1H). We found that, compared to *N*-RAM^-^ neurons, *N*-RAM^+^ neurons displayed significantly larger oIPSC amplitudes upon stimulation of CCK^+^ interneurons, with no significant differences observed upon stimulation of PV^+^ or SST^+^ interneurons (Figure 1I-J). In contrast, oIPSC amplitudes on *F*-RAM^+^ and *F*-RAM^-^ neurons were comparable (Figure 1I-J). Therefore, *N*-RAM^+^ but not *F*-RAM^+^ neurons exhibit inhibitory synaptic plasticity by strengthening their inhibitory input selectively from CCK^+^ interneurons.

Together, our results indicated that *Fos*- and *Npas4* -dependent ensembles within an engram in the mouse DG exhibit synaptic plasticity in a transcription-specific manner: *Fos*-dependent ensembles exhibit excitatory plasticity in feedforward synapses from the MEC, while *Npas4* -dependent ensembles exhibit inhibitory plasticity in recurrent synapses from CCK^+^ interneurons.

### Transcription-specific synaptic plasticity shapes engram stability

We proposed a computational model to investigate the link between transcription-specific synaptic plasticity and the computations promoted by *Fos*- and *Npas4* -dependent ensembles in the DG (Figure 2A). Specifically, our spiking neural network model consisted of an input population that projected to a DG network, where excitatory neurons modeled granule cells and inhibitory neurons modeled CCK^+^ interneurons. Upon initialization, the network model underwent a burn-in period to stabilize neural activity. During this period, we assumed that the entire population of excitatory neurons expressed both excitatory and inhibitory forms of long-term synaptic plasticity, in line with their role in the stabilization of neural activity during development.^29^ Long-term excitatory synaptic plasticity combined Hebbian terms, including long-term potentiation (LTP) and long-term depression (LTD) in the form of triplet spike-timing-dependent plasticity (STDP),^30^ and non-Hebbian terms, consisting of heterosynaptic plasticity^31^ and transmitter-induced plasticity,^32^ as previously proposed.^33^ Triplet STDP enabled Hebbian correlation-based learning. Heterosynaptic plasticity prevented excessive LTP of excitatory synapses and an ensuing pathological increase in neural activity. Transmitter-induced plasticity prevented neurons from becoming silent. Previous work showed that this combination of Hebbian and non-Hebbian forms of excitatory plasticity can support stable memory encoding and recall.^33^ Inhibitory synaptic plasticity took the form of homeostatic STDP, including Hebbian, anti-Hebbian, and non-Hebbian terms that acted to maintain excitatory neural activity at a set target level as previously proposed.^33, 34^ At the end of the burn-in period, we assumed that half of the population of excitatory neurons expressed exclusively *Fos*-dependent excitatory synaptic plasticity (*Fos*^+^ neurons), while the other half expressed exclusively *Npas4* -dependent inhibitory synaptic plasticity (*Npas4* ^+^ neurons). This was consistent with both (1) our previous work showing that *Fos*- and *Npas4* -dependent ensembles in the mouse DG are distinct neuronal subpopulations^17^ and (2) our experiments showing that they express distinct forms of synaptic plasticity (Figure 1E/J). Importantly, *Fos*-dependent excitatory synaptic plasticity included only Hebbian LTP and heterosynaptic plasticity, while *Npas4* -dependent inhibitory synaptic plasticity included only a Hebbian term (see Methods for a detailed description of the model).

**Figure 2.**
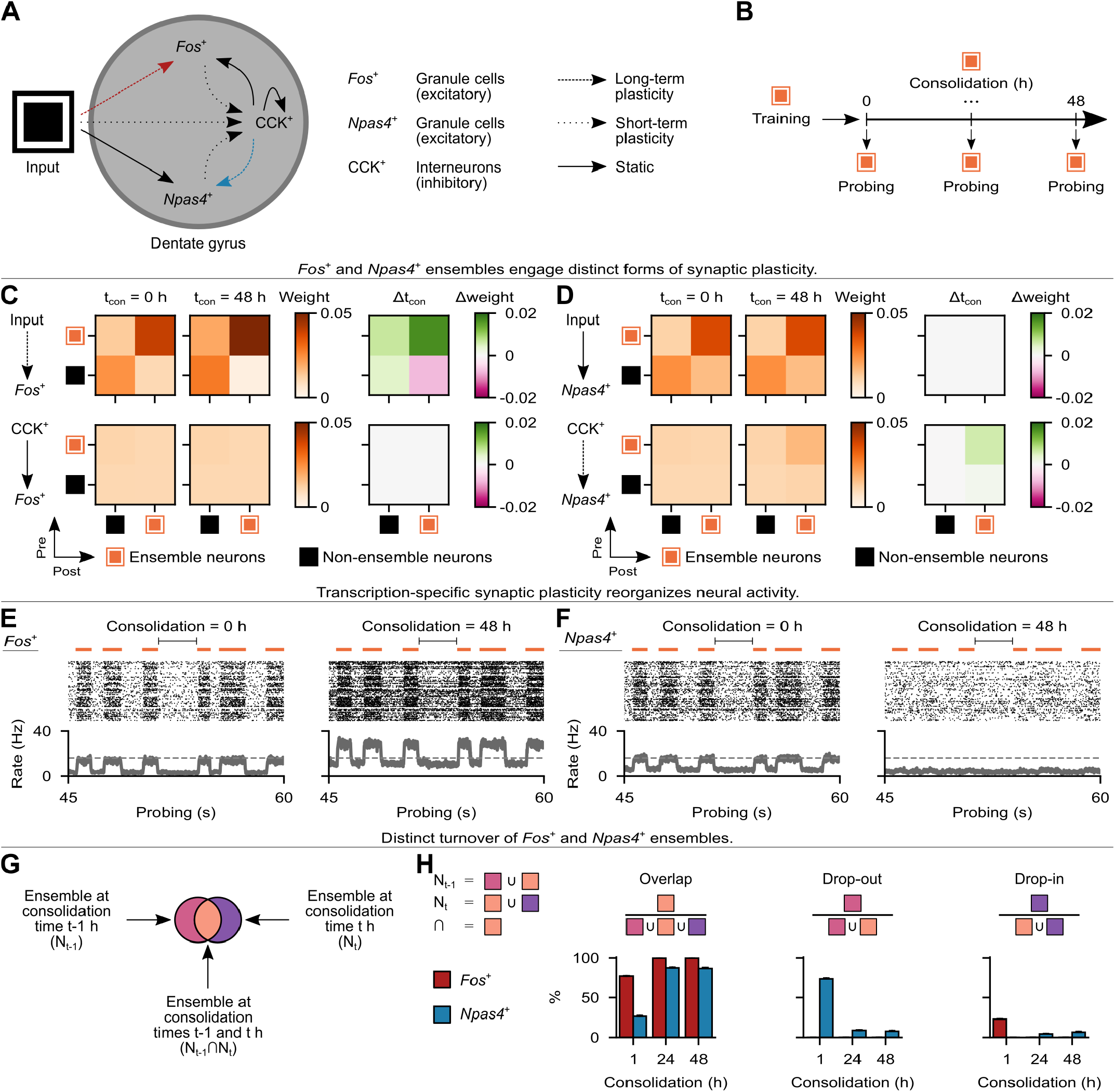
Transcription-specific synaptic plasticity reshapes neuronal ensembles within an engram. (A) Schematic of the computational model. Left: input population and DG network with *Fos*^+^ and *Npas4* ^+^ granule cells (excitatory) as well as CCK^+^ interneurons (inhibitory). Right: plasticity of feedforward and recurrent synapses (see Methods). (B) Schematic of the simulation protocol with probing periods (see Methods). (C-D) Mean weight strength of synapses onto *Fos*^+^ (C) or *Npas4* ^+^ (D) neurons clustered according to ensemble status (i.e., ensemble and non-ensemble neurons) at the end of training (i.e., 0 h of consolidation). Top: feedforward excitatory synapses. Bottom: recurrent inhibitory synapses. Left: at 0 h of consolidation. Middle: at 48 h of consolidation. Right: change in weight strength between 0 and 48 h of consolidation. Plasticity of synapses as in (A). Representative trial shown. (E-F) Activity of *Fos*^+^ (E) or *Npas4* ^+^ (F) neurons during probing at 0 h (left) and 48 h (right) of consolidation. Top: training stimulus presentation times (in orange). Middle: spike raster of 256 randomly-chosen neurons (for clarity, only every fifth spike is shown). Bottom: average firing rate of the population of neurons (dashed line indicates the activation threshold *ζ*^*thr*^ = 16 Hz). Population rate shown without smoothing or convolution. Representative trial shown. (G) Schematic of neuronal ensembles over the course of consolidation. Each circle denotes the set of ensemble neurons at a given time point. (H) Post-encoding evolution of neuronal ensembles. Overlap (left), drop-out (middle), and drop-in (right) measured between consecutive time points as in (G). Data shown as mean ± SEM. *n* = 10 trials.

After the burn-in period, our network followed a memory acquisition and consolidation protocol^35^ (Figure 2B). Specifically, the network was subjected to a training period when a training stimulus was presented to simulate an associative learning task. Next, the network proceeded to a consolidation period when the training stimulus was reactivated, consistent with previous reports that training-activated neurons in the entorhinal cortex of rodents and humans are selectively reactivated during post-training periods of sleep or rest.^36, 37^ At regular intervals throughout the consolidation period (consolidation = 0, 1, …, 48 h), the network advanced to a probing period to identify neuronal ensembles encoding the training stimulus at a given time point. In particular, we identified *Fos*^+^, *Npas4* ^+^, and CCK^+^ ensembles by examining which cells were selectively activated in response to the training stimulus during probing (i.e., average stimulus-evoked firing rate above the activation threshold *ζ*^*thr*^ = 16 Hz). Training stimulus neurons in the input population were considered a fixed ensemble (see Methods for a detailed description of neuronal ensembles in our simulations).

We found that feedforward excitatory synapses from input ensemble neurons onto *Fos*^+^ ensemble neurons were strengthened, while synapses from input non-ensemble neurons onto *Fos*^+^ ensemble neurons were weakened, in line with previous experiments showing cross-region synaptic coupling between ensemble neurons^38^ (Figure 2C). Recurrent inhibitory synapses onto *Fos*^+^ neurons remained static. On the other hand, feedforward excitatory synapses onto *Npas4* ^+^ neurons remained static, while recurrent inhibitory synapses from CCK^+^ ensemble interneurons onto *Npas4* ^+^ ensemble neurons were strengthened (Figure 2D). This was consistent with our experiments showing that *Fos*- and *Npas4* -dependent ensembles express excitatory and inhibitory synaptic plasticity, respectively (Figure 1E/J). Note that the weight strength of synapses onto *Fos* ^+^ or *Npas4* ^+^ neurons in the network model stabilized over the course of consolidation (Figure S1A-B). Furthermore, the population of *Fos*^+^ neurons increased its baseline firing rate and its training stimulus-evoked response with consolidation (Figure 2E). Conversely, *Npas4* ^+^ neurons nearly stopped responding to the training stimulus at the population level as consolidation progressed (Figure 2F). These opposing effects on the population responses of *Fos*^+^ and *Npas4*^+^ neurons could be attributed to the distinct forms of synaptic plasticity they express: *Fos*-dependent plasticity potentiated excitatory synapses onto *Fos*^+^ neurons without a compensatory potentiation of inhibition, whereas *Npas4* -dependent plasticity potentiated inhibitory synapses onto *Npas4*^+^ neurons to maintain their activity at a homeostatic target level. Despite these differences, the activity of both *Fos*^+^ and *Npas4*^+^ neurons remained stable (Figure S1C-D). CCK^+^ interneurons also maintained stable activity (Figure S2A-B).

We then examined the temporal evolution of neuronal ensembles in our simulations by computing the overlap, drop-out, and drop-in between consecutive consolidation time points (Figure 2G-H). *Fos*^+^ ensembles had a relatively high overlap in the initial periods of consolidation, with only a moderate drop-in and no drop-out. In the later stages of consolidation, their overlap increased further with the decay of drop-in. In contrast, *Npas4*^+^ ensembles initially had a low overlap accompanied by a high drop-out but no drop-in. Later, their overlap increased substantially, with a considerable decrease in drop-out and only a modest increase in drop-in. CCK^+^ ensembles remained stable (i.e., without turnover) (Figure S2C). Thus, our model suggests that, following initial memory encoding, *Fos*- and *Npas4* -dependent ensembles exhibit drop-in and drop-out, respectively, but stabilize as consolidation progresses. This is in line with our previous work showing that neuronal ensembles in mouse DG exhibit high rates of drop-in and drop-out following CFC.^35^

### Transcription-specific synaptic plasticity enables engram computations

To evaluate the computational role of *Fos*^+^ and *Npas4*^+^ ensembles in our model, we used a modified memory acquisition and recall protocol^35^ (Figure 3A). Specifically, our network model was initially subjected to a training period and then proceeded to a consolidation period as in our previous protocol (compare Figure 3A to Figure 2B). However, at regular intervals throughout the consolidation period (consolidation = 0, 1, …, 48 h), the network advanced to a recall period when partial cues of the training stimulus or a novel, unseen stimulus were presented (Figure 3B). Note that there were a total of four separate recall periods (i.e., one for the training stimulus and one for each of the three novel stimuli) after every sampled consolidation interval. The intersection between the training stimulus (i.e., square) and each novel stimulus (i.e., circle, hexagon, and pentagon) varied roughly from 50 to 60% (Figure 3C). At the end of training (i.e., 0 h of consolidation), we found that *Fos*^+^ ensembles on average fired above the activation threshold *ζ*^*thr*^ = 16 Hz in response to cues of the training stimulus but not the novel stimuli, while *Npas4*^+^ ensembles on average fired above the activation threshold in response to cues of both the training stimulus and the novel stimuli (Figure 3D, top row). As consolidation progressed, these patterns reversed: *Fos*^+^ ensembles on average responded above the activation threshold to cues of both the training stimulus and the novel stimuli, whereas *Npas4*^+^ ensembles on average responded above the activation threshold to cues of the training stimulus but not the novel stimuli. We considered an ensemble activated by a partial cue when its cue-evoked firing rate exceeded the activation threshold. Ensemble activation thus served as a proxy for recall. Under this criterion, *Fos*^+^ ensembles were initially activated more often by cues of the training stimulus than the novel stimuli, but this difference nearly vanished with consolidation (Figure 3D, middle left panel). In contrast, *Npas4*^+^ ensembles were initially activated equally by cues of the training and novel stimuli, but became preferentially activated by cues of the training stimulus over the course of consolidation (Figure 3D, middle right panel).

**Figure 3.**
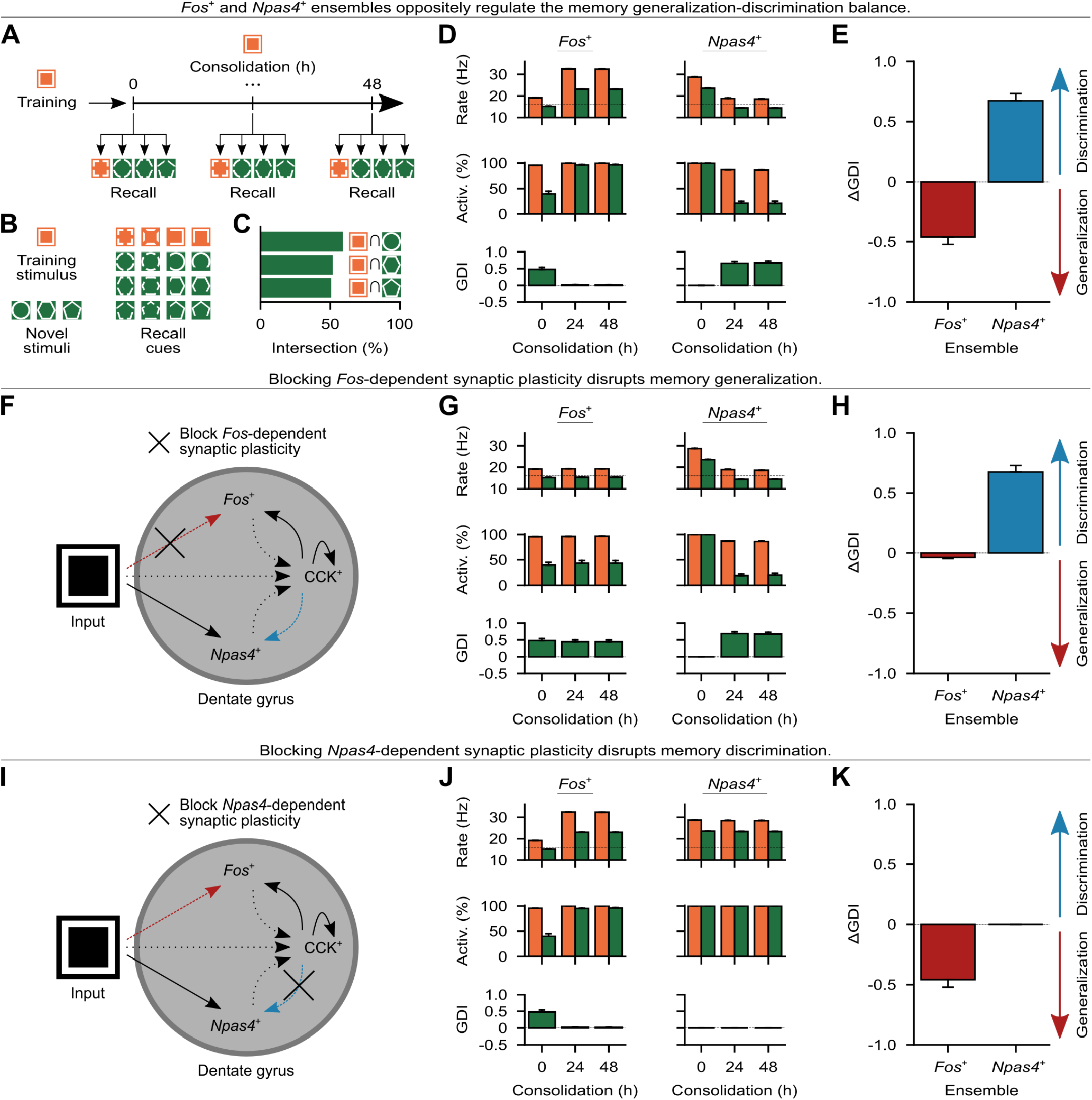
Transcription-specific synaptic plasticity drives ensemble computations. (A) Schematic of the simulation protocol with recall periods (see Methods). (B) Schematic of the training and novel stimuli with associated partial cues for recall. (C) Intersection between the training stimulus and each novel stimulus as a fraction of training stimulus neurons. (D) Top: cue-evoked ensemble firing rate (dashed line indicates the activation threshold *ζ*^*thr*^ = 16 Hz). Middle: cue-evoked ensemble activation. Bottom: GDI between the ensemble activation evoked by cues of the training and novel stimuli. Left: *Fos*^+^ ensemble. Right: *Npas4*^+^ ensemble. Color denotes stimulus as in (B). (E) Change in ensemble activation GDI between 0 and 48 h of consolidation. A decrease in GDI indicates generalization, whereas an increase indicates discrimination. (F) Schematic of the computational model with *Fos*-dependent excitatory synaptic plasticity blocked during consolidation (compare to Figure 2A). (G-H) Same as (D-E) but when the model in (F) was subjected to the simulation protocol in (A). (I) Schematic of the computational model with *Npas4* -dependent inhibitory synaptic plasticity blocked during consolidation (compare to Figure 2A). (J-K) Same as (D-E) but when the model in (I) was subjected to the simulation protocol in (A). Data shown as mean ± SEM and *n* = 10 trials in (D-E), (G-H), and (J-K).

We computed a generalization-discrimination index (GDI) that measured the normalized difference between the ensemble activation evoked by cues of the training stimulus and the novel stimuli. We found that *Fos*^+^ and *Npas4*^+^ ensembles changed their activation GDI in opposite directions, with *Fos*^+^ ensembles decreasing and *Npas4*^+^ ensembles increasing their GDI with consolidation (Figure 3D, bottom row). We defined memory generalization and discrimination at the neural level as a decrease or an increase in ensemble activation GDI, respectively. Intuitively, we considered that neuronal ensembles promoted generalization when they were preferentially activated by the training stimulus at the end of training (i.e., when ensembles emerged), but became activated equally by the training and novel stimuli with consolidation. Conversely, we considered that neuronal ensembles promoted discrimination when they were activated equally by the training and novel stimuli at the end of training, but became preferentially activated by the training stimulus with consolidation. Under this framework, *Fos*^+^ and *Npas4*^+^ ensembles promoted generalization and discrimination, respectively (Figure 3E). This was consistent with our previous work on the functional role of *Fos*- and *Npas4* -dependent ensembles in mouse DG.^17^ Note that CCK^+^ ensembles in our network model failed to promote generalization or discrimination since their activation GDI did not change over the course of consolidation (Figure S2D).

We conducted a sensitivity analysis of the activation threshold to examine its effect on ensemble computations (Figure S2E). We found that there was a range of threshold values (i.e., 14–18 Hz) for which *Fos*^+^ and *Npas4*^+^ ensembles promoted generalization (i.e., ΔGDI *<* 0) and discrimination (i.e., ΔGDI > 0), respectively, while maintaining their training stimulus cue-evoked activation above 70%. We also explored how changing our model of transcription-specific synaptic plasticity affects ensemble computations. In particular, removing Hebbian LTP or the heterosynaptic term from *Fos*-dependent excitatory plasticity prevented *Fos*^+^ ensembles from promoting generalization (i.e., ΔGDI = 0) (Figure S3A-F). On the other hand, adding Hebbian LTD or the transmitter-induced term to *Fos*-dependent excitatory plasticity did not prevent *Fos*^+^ ensembles from promoting generalization (i.e., ΔGDI *<* 0) (Figure S3G-L). Note, however, that adding Hebbian LTD substantially impaired generalization (i.e., ΔGDI less negative than in control). Interestingly, including the anti-Hebbian but not the non-Hebbian term in *Npas4* -dependent inhibitory plasticity kept *Npas4*^+^ ensembles promoting discrimination (i.e., ΔGDI > 0) (Figure S4). Together, these results showed that multiple forms of *Fos*- and *Npas4* -dependent synaptic plasticity can support generalization and discrimination, respectively, pointing to functionally degenerate forms of transcription-specific plasticity.

Next, we used our model to investigate whether transcription-specific synaptic plasticity has a causal role in ensemble computations. We found that blocking *Fos*-dependent excitatory synaptic plasticity in effect prevented *Fos*^+^ ensembles from decreasing their activation GDI with consolidation, while *Npas4*^+^ ensembles reliably increased their activation GDI (Figure 3F-G). Consequently, *Fos*^+^ ensembles effectively failed to promote generalization (i.e., ΔGDI ∼0), whereas *Npas4*^+^ ensembles continued to promote discrimination (i.e., ΔGDI > 0) (Figure 3H). In contrast, blocking *Npas4* -dependent inhibitory synaptic plasticity prevented *Npas4*^+^ ensembles from increasing their activation GDI with consolidation, while *Fos*^+^ ensembles reliably decreased their activation GDI (Figure 3I-J). As a result, *Npas4*^+^ ensembles failed to promote discrimination (i.e., ΔGDI = 0), whereas *Fos*^+^ ensembles continued to promote generalization (i.e., ΔGDI *<* 0) (Figure 3K). Importantly, both *Fos*^+^ and *Npas4*^+^ ensembles remained highly activated by cues of the training stimulus when either *Fos*- or *Npas4* -dependent synaptic plasticity was blocked, indicating that neither manipulation impaired memory recall (Figure 3G/J, middle rows). Thus, our model predicted that blocking *Fos*- or *Npas4* -dependent synaptic plasticity disrupts generalization and discrimination, respectively.

### Transcription-specific synaptic plasticity mediates contextual fear generalization and discrimination

To test our model’s prediction that transcription-specific synaptic plasticity has a causal role in generalization and discrimination, we sought to block *Fos*- and *Npas4* -dependent synaptic plasticity *in vivo*. Although FOS and NPAS4 are widely used as markers of neuronal activation,^11, 39^ it remains unknown whether they regulate the downstream transcriptional programs that instruct the specific synaptic changes associated with the active neuronal ensembles they define. Given that *Fos*- and *Npas4* -dependent ensembles exhibit distinct forms of synaptic plasticity (Figure 1E/J), we asked whether *Fos* and *Npas4* mediate their expression.

To determine whether *Fos* regulates learning-induced excitatory synaptic plasticity, we knocked out *Fos* using CRISPR by co-injecting AAVs expressing Cas9 and *Fos*-targeting gRNA into the DG (Figure 4A, left panel). We verified knockout efficiency by administering the glutamate receptor agonist kainic acid (KA) via intraperitoneal (i.p.) injection to induce seizure, depolarizing hippocampal neurons. Subsequent immunostaining revealed a robust reduction in FOS protein expression in the *Fos* gRNA group compared to scramble gRNA controls, confirming highly efficient gene deletion *in vivo* (Figure 4A, right panel). To label the *Fos*-dependent ensemble, we used a *Fos*-promoter-driven reporter (mKate2) under Dox-dependent Tet-off control (Figure 4B). Following CFC, we recorded miniature excitatory postsynaptic currents (mEPSCs) from *Fos*^+^ and *Fos*^-^ neighboring neurons in the DG (Figure 4C). In control mice, *Fos*^+^ neurons exhibited a significantly higher mEPSC frequency compared to neighboring *Fos*^-^ neurons. This difference vanished in *Fos* knockout mice, indicating that the observed learning-induced excitatory synaptic enhancement was effectively abolished by *Fos* deletion.

**Figure 4.**
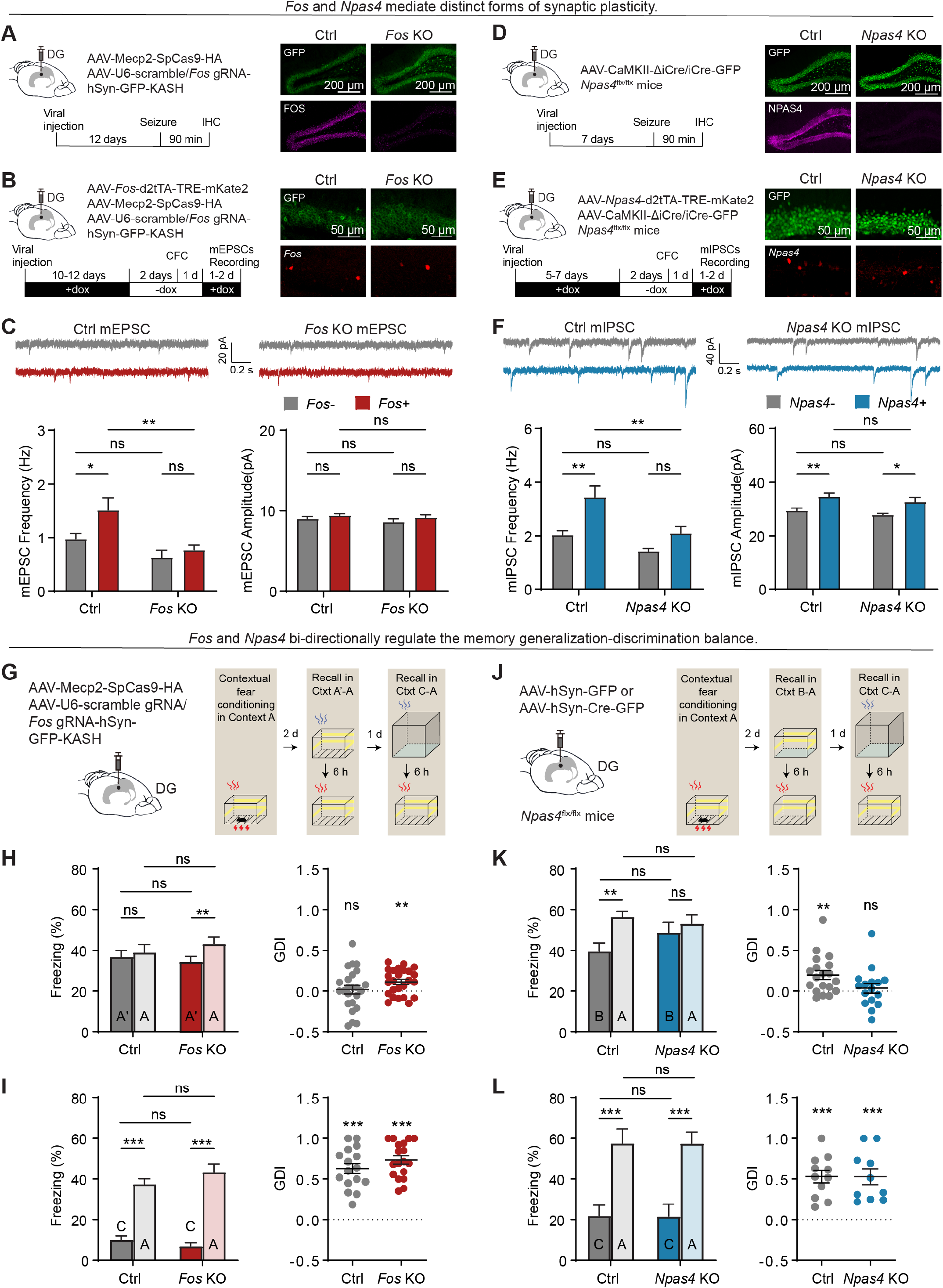
*Fos*- and *Npas4* -dependent synaptic plasticity bi-directionally regulate the memory generalization-discrimination balance. (A) Validation of *Fos* conditional knockout. Left: schematic illustration of the viral strategy for *Fos* deletion. Right: representative images showing *Fos* knockout efficiency. (B) Experimental strategy for *Fos* ensemble labeling and mEPSC recording. Left: schematic illustration of the viral strategy for *Fos* conditional knockout and *Fos* ensemble labeling. Right: representative images showing the labeling of *Fos*^+^ neurons for mEPSC recording. (C) Top: representative mEPSC traces on pairs of *Fos*^+^ and *Fos*^-^ nearby neurons under control (left) and *Fos* knockout (right) conditions. Bottom: mEPSC frequency (left) and amplitude (right) on pairs of *Fos*^+^ and *Fos*^-^ nearby neurons under control and *Fos* knockout conditions (*n* = 13 neurons per group). (D) Validation of *Npas4* conditional knockout. Left: schematic illustration of the viral strategy for *Npas4* deletion. Right: representative images showing *Npas4* knockout efficiency. (E) Experimental strategy for *Npas4* ensemble labeling and mIPSC recording. Left: schematic illustration of the viral strategy for *Npas4* conditional knockout and *Npas4* ensemble labeling. Right: representative images showing the labeling of *Npas4*^+^ neurons for mIPSC recording. (F) Top: representative mIPSC traces on pairs of *Npas4*^+^ and *Npas4*^-^ nearby neurons under control (left) and *Npas4* knockout (right) conditions. Bottom: mIPSC frequency (left) and amplitude (right) on pairs of *Npas4*^+^ and *Npas4*^-^ nearby neurons under control and *Npas4* knockout conditions (*n* = 14 neurons per group). (G) Experimental scheme for *Fos* knockout and associated behavioral assay to test memory generalization-discrimination balance. Left: schematic illustration of the viral strategy used to knock out *Fos* in the DG. Right: experimental timeline of the contextual fear memory generalization-discrimination test. Mice were initially fear-conditioned in context A. Following a 48-h post-conditioning period, mice were subjected to recall sessions in (1) a highly similar context A’ followed by the training context A on day 1, and (2) a highly distinct context C followed by the training context A on day 2 (see Methods). (H) Freezing levels (left) and corresponding GDI (right) for recall in context A’ and A under control (*n* = 24 mice) and *Fos* knockout (*n* = 25 mice) conditions. (I) Freezing levels (left) and corresponding GDI (right) for recall in context C and A under control (*n* = 16 mice) and *Fos* knockout (*n* = 18 mice) conditions. (J) Experimental scheme for *Npas4* knockout and associated behavioral assay to test memory generalization-discrimination balance. Left: schematic illustration of the viral strategy used to knock out *Npas4* in the DG. Right: experimental timeline of the contextual fear memory generalization-discrimination test. Mice were initially fear-conditioned in context A. Following a 48-h post-conditioning period, mice were subjected to recall sessions in (1) a moderately similar context B followed by the training context A on day 1, and (2) a highly distinct context C followed by the training context A on day 2 (see Methods). (K) Freezing levels (left) and corresponding GDI (right) for recall in context B and A under control (*n* = 19 mice) and *Npas4* knockout (*n* = 16 mice) conditions. (L) Freezing levels (left) and corresponding GDI (right) for recall in context C and A under control (*n* = 11 mice) and *Npas4* knockout (*n* = 10 mice) conditions. Two-way mixed ANOVA with Sidak’s multiple comparisons test was used for mEPSCs, mIPSCs, and freezing levels. One-sample *t* test was used to compare freezing behavior GDI to zero. Data shown as mean ± SEM. \**p <* 0.05, \*\**p <* 0.01, \*\*\**p <* 0.001.

To test whether *Npas4* regulates learning-induced inhibitory synaptic plasticity, we knocked out *Npas4* by injecting an AAV expressing Cre recombinase into the DG of *Npas4* conditional knockout mice (Figure 4D, left panel). The knockout efficiency was confirmed by KA-induced seizure followed by immunostaining of NPAS4, which revealed a robust reduction in NPAS4 protein expression in the Cre group compared to inactive delta Cre controls (Figure 4D, right panel). We labeled the *Npas4* - dependent ensemble using a minimal *Npas4* -promoter-driven reporter (mKate2) under Tet-off control (Figure 4E). Following CFC, we recorded miniature inhibitory postsynaptic currents (mIPSCs) from *Npas4*^+^ and *Npas4*^-^ neighboring neurons in the DG (Figure 4F). In control mice, *Npas4*^+^ neurons exhibited a significantly higher mIPSC frequency compared to neighboring *Npas4*^-^ neurons. This difference was eliminated in *Npas4* knockout mice, indicating that the observed learning-induced inhibitory synaptic enhancement was effectively abolished by *Npas4* deletion. Please note that *Npas4*^+^ neurons exhibited larger mIPSC amplitudes compared to *Npas4*^-^ neurons under both control and knockout conditions, suggesting that this effect is independent of *Npas4*. Together, these findings demonstrated that *Fos* is crucial for learning-induced enhancement of excitatory synaptic inputs onto *Fos*-dependent ensembles, whereas *Npas4* is essential for the selective recruitment of inhibitory synaptic inputs onto *Npas4* -dependent ensembles.

Next, we used our *Fos* and *Npas4* knockout strategies to test our model’s prediction that transcription-specific synaptic plasticity mediates ensemble computations. Specifically, we acutely knocked out *Fos* in the DG of mice that were later subjected to a contextual fear memory generalization-discrimination assay (Figure 4G). Following fear conditioning in context A, mice were tested in a series of recall sessions conducted in contexts with varying degrees of similarity to the training context A. On the first recall day, mice were exposed to a highly similar context A’ followed by exposure to the training context A. On the following day, mice were exposed to a highly distinct context C and then to the training context A. We found that control mice exhibited comparable freezing levels in context A and the highly similar context A’, while *Fos* knockout mice showed significantly lower freezing levels in context A’ compared to context A (Figure 4H, left panel). To evaluate whether mice behaviorally generalized or discriminated, we computed a freezing behavior GDI that measured the normalized difference between freezing levels in the training context and a novel context, in a manner analogous to the ensemble activation GDI previously used to characterize generalization and discrimination at the neural level. When freezing behavior GDI was zero (i.e., comparable freezing levels in the training and novel contexts), mice were considered to generalize (or fail to discriminate); when it was different from zero (i.e., different freezing levels in the training and novel contexts), they were considered to discriminate (or fail to generalize). Under this criterion, control mice generalized between contexts A and A’, while *Fos* knockout mice failed to do so (Figure 4H, right panel). In contrast, both control and *Fos* knockout mice displayed higher freezing levels in context A than in the highly distinct context C, indicating that the two groups were able to discriminate between these two contexts (Figure 4I). Thus, knocking out *Fos* disrupted generalization of highly similar contexts, but not discrimination of highly distinct ones.

Furthermore, we acutely knocked out *Npas4* in the DG of mice that were subsequently subjected to a contextual fear memory generalization-discrimination assay analogous to the one previously used in our *Fos* knockout experiments (Figure 4J, compare to Figure 4G). Following fear conditioning in context A, mice were tested in recall sessions conducted in contexts with varying degrees of similarity to the training context A. On the first recall day, mice were placed in a moderately similar context B and subsequently in the training context A. On the following day, mice were placed in a highly distinct context C and the training context A. We found that control mice exhibited higher freezing levels in context A than in the moderately similar context B, while this difference was eliminated in *Npas4* knockout mice (Figure 4K, left panel). Freezing behavior GDI showed that control mice discriminated between contexts A and B, while *Npas4* knockout mice failed to do so (Figure 4K, right panel). Notably, both control and *Npas4* knockout mice were able to discriminate between context A and the highly distinct context C (Figure 4L). Hence, knocking out *Npas4* disrupted discrimination of moderately similar contexts, but not of highly distinct ones. Consistent with our model, neither *Fos* nor *Npas4* knockout impaired memory recall, as evidenced by the comparable freezing levels in the training context A under control and knockout conditions (Figure 4H-I/K-L, left panels).

Together, our results supported our model’s prediction that transcription-specific synaptic plasticity has a causal role in engram computations, with *Fos*-dependent excitatory synaptic plasticity and *Npas4* -dependent inhibitory synaptic plasticity mediating generalization and discrimination, respectively.

## Discussion

Here, we provided computational and experimental evidence that transcription-specific synaptic plasticity regulates memory computations. Informed by our electrophysiology experiments, we proposed a spiking neural network model of the DG incorporating *Fos*-dependent excitatory synaptic plasticity and *Npas4* -dependent inhibitory synaptic plasticity. Our model predicted that blocking *Fos*- or *Npas4* -dependent synaptic plasticity disrupts memory generalization and discrimination, respectively, without impairing recall. To test this prediction, we developed *Fos* and *Npas4* knockout strategies that selectively targeted the expression of synaptic plasticity in a transcription-specific manner. Associative learning experiments with acute *Fos* or *Npas4* knockout in the DG supported our model’s prediction. Together, our converging *in silico* and *in vivo* findings established a causal link between transcriptional regulation of synaptic plasticity and computational versatility in an engram.

Previous studies showed that activity-dependent transcription and synaptic plasticity are crucial for memory formation and recall.^6–8, 15, 16^ Critically, whereas our previous work found that an individual memory engram contains transcriptionally defined neuronal ensembles that exhibit specific forms of synaptic plasticity and are functionally distinct,^17^ our present study revealed that transcription-specific synaptic plasticity itself regulates ensemble computations. In particular, we showed that blocking synaptic plasticity in a transcription-specific manner (i.e., *Fos*- or *Npas4* -dependent plasticity) disrupts a specific ensemble computation (i.e., generalization or discrimination), but not memory recall. By linking heterogeneous forms of transcriptionally regulated plasticity within the same engram to distinct ensemble computations, our findings extended the role of synaptic plasticity beyond memory storage to the active control of memory computations.

While our work focused on associative memory in the DG, transcription-specific synaptic plasticity may have a fundamental computational role in sensory, cognitive, and motor functions throughout the brain. Hundreds of transcriptomic cell types have been identified across the human, chimpanzee, gorilla, macaque, and mouse brains,^40–46^ and a growing body of evidence has linked activity-dependent transcription to synaptic plasticity in perception, learning, and memory.^11, 47^ This raises the possibility that neural representations in general (e.g., sensory, associative, spatial, episodic, semantic, and motor) may engage transcription-specific synaptic plasticity to mediate particular computations, thereby providing a neural substrate for adaptive, experience-guided behavior. Importantly, our work causally linked *Fos*-dependent excitatory synaptic plasticity to generalization and *Npas4* -dependent inhibitory synaptic plasticity to discrimination, illustrating how linking activity-dependent transcription, synaptic plasticity, and computation can uncover the connection between transcriptomic heterogeneity and functional diversity in neural circuits.

Our spiking neural network model with transcription-specific synaptic plasticity highlighted the computational benefits of incorporating heterogeneity at the level of plasticity. Previous models with stable memory encoding and recall included multiple co-active forms of synaptic plasticity on the same neuronal population (i.e., excitatory or inhibitory) that were homogeneously engaged (e.g., all excitatory neurons engaged the same form of Hebbian plasticity in their recurrent excitatory synapses and the same form of homeostatic plasticity in their recurrent inhibitory synapses).^33, 48, 49^ In contrast, our model, inspired by our experimental findings, incorporated transcription-specific synaptic plasticity by having half of the population of excitatory neurons engage plasticity only in their feedforward excitatory synapses (*Fos*^+^ neurons) and the other half engage plasticity only in their recurrent inhibitory synapses (*Npas4*^+^ neurons). This enabled the emergence, within the same population, of distinct neuronal ensembles that promoted opposite and complementary computations (i.e., generalization and discrimination). Notably, recently proposed network models have begun to incorporate heterogeneous expression of synaptic plasticity by endowing synapses with broadly distributed learning rates^50^ or the ability to independently switch between two different learning rules.^51^ Our work serves as a proof of principle that including heterogeneity in the form of transcription-specific synaptic plasticity can enrich the computational repertoire of neural networks.

Our computational model also provided insights into the temporal evolution and computational strategies of *Fos*- and *Npas4*-dependent ensembles. In our simulations, *Fos*-dependent synaptic plasticity recruited new neurons into *Fos*^+^ ensembles by potentiating excitatory synapses onto *Fos*^+^ neurons that were initially unresponsive to the training stimulus. In contrast, *Npas4*-dependent synaptic plasticity forced neurons out of *Npas4*^+^ ensembles by potentiating inhibitory synapses onto *Npas4*^+^ neurons that were initially responsive to the training stimulus. Consequently, *Fos*^+^ and *Npas4*^+^ ensembles exhibited drop-in and drop-out, respectively. Furthermore, *Fos*^+^ ensembles promoted generalization by selectively increasing the training stimulus-evoked response of *Fos*^+^ neurons, with new neurons dropping into the ensemble expanding the set of feedforward synaptic inputs that could trigger an ensemble response. Conversely, *Npas4*^+^ ensembles promoted discrimination by selectively decreasing the training stimulus-evoked response of *Npas4*^+^ neurons, with neurons dropping out of the ensemble driving sparsification and competition such that only the most responsive neurons could remain part of the ensemble. Whereas our prior work associated neuronal ensemble turnover with discrimination,^35^ our present study specifically linked *Fos*^+^ ensemble drop-in and *Npas4*^+^ ensemble drop-out with generalization and discrimination, respectively. Importantly, expressing *Fos*- and *Npas4* -dependent synaptic plasticity in distinct neuronal subpopulations allowed our network model to make both generalization and discrimination available upon presentation of a single novel stimulus similar to the training stimulus: the network output could be “read out” from either *Fos*^+^ or *Npas4*^+^ ensembles. By comparison, previous network models could only generalize or discriminate when presented with a given novel stimulus, as homogeneous expression of synaptic plasticity led to the encoding of a single neuronal ensemble per training stimulus.^33, 35, 48, 49^ Thus, our model suggests that the organization of an engram into distinct neuronal ensembles that engage transcriptionally regulated forms of plasticity orthogonalizes neural computations and thereby increases cognitive flexibility.

Together, our computational and experimental results revealed that the transcriptional regulation of synaptic plasticity exerts fine control over memory computations. In particular, we showed that transcription-specific synaptic plasticity within an engram mediates the generalization-discrimination balance. This has implications for both adaptive memory computations (e.g., selective memory expression) and their maladaptive counterparts (e.g., overgeneralized fear expression in PTSD). Future work will be needed to investigate the link between transcription, synaptic plasticity, and computation across functional domains and brain regions.

## Data and code availability

The data generated by our experiments, together with our simulation and data analysis code, will be made publicly available upon publication.

## Acknowledgments

We thank Andrea Navas-Olive for feedback on an earlier version of this manuscript. This research was funded in whole or in part by the Austrian Science Fund (FWF) 10.55776/COE16 to D.F.T. This project has received funding from the HORIZON EUROPE European Research Council (ERC) consolidator grant SYNAPSEEK to T.P.V. This work was supported by NIH grants NS115543, DC014701, and MH116673 to Y.L. This work was supported by Wellcome Trust 200790/Z/16/Z, EPSRC EP/R035806/1, and ERC MotorAdapt 101169605 to C.C.

## Author contributions

D.F.T. conceptualized the model, performed the simulations, and analyzed the simulation output. Y.L., M.M., and X.S. designed the experiments. M.M. performed or participated in all experiments. X.S. contributed to the CRISPR gRNA design and behavior. L.Y. contributed to electrophysiological experiments. T.P.V., Y.L., and C.C. supervised the project. D.F.T. and M.M. wrote the original draft. All authors reviewed and edited the final manuscript.

## Methods

### Subjects

8–12-week-old male C57BL/6 mice were used in all experiments. Wild-type mice were purchased from Charles River Laboratories. Npas4^flx/flx^ (*Npas4* conditional knockout) mice were generated previously.^52^ PV-Cre (Pvalb^tm1(cre)Arbr^/J), CCK-Cre (Cck^tm1.1(cre)Zjh^/J), and SST-Cre (Sst^tm2.1(cre)Zjh^/J) mice were purchased from Jackson Laboratory. Heterozygous mice carrying the Cre allele were bred with wild-type animals from Charles River Laboratories for more than three generations. All mice were housed with a 12-hour light-dark cycle. Animal protocols were performed in accordance with the US National Institutes of Health (NIH) guidelines and approved by the Institutional Animal Care and Use Committee at University of Texas Southwestern Medical Center.

### Viral vectors

All AAVs were produced in house using a previously described protocol.^53^ Viral dilutions were determined for individual experiments using pilot injections.

In experiments to label *F*-RAM^+^ or *N*-RAM^+^ ensembles, AAV9-*F*-RAM-d2tTA-TRE-mKate2 and AAV9-*N*-RAM-d2tTA-TRE-mKate2 viruses were used, respectively. To test the plasticity of excitatory inputs, AAV9-CaMKII-ChR2-EYFP was injected into LEC, MEC or contralateral DG. To test the plasticity of inhibitory inputs, AAV9-Dlx5/6-DIO-ChR2-EYFP was injected into CCK-, PV-, or SST-Cre mice.

To test whether *Fos* mediates the plasticity of excitatory inputs, AAV9-U6-*Fos* gRNA-hSyn-GFP-KASH or AAV9-U6-scramble gRNA-hSyn-GFP-KASH (hSyn-GFP-KASH was used to determine injection accuracy and infection efficiency) were co-injected with AAV9-Mecp2-Cas9-HA and AAV9-*Fos*-dt2tTA-TRE-mKate2. To test whether *Npas4* mediates the plasticity of inhibitory inputs, AAV9-CaMKII-iCre-GFP or AAV9-CaMKII-delta iCre-GFP were co-injected with AAV9-*Npas4* -dt2tTA-TRE-mKate2 into *Npas4* conditional knockout mice.

To determine whether *Fos* regulates the memory generalization-discrimination balance, AAV9-U6-*Fos* gRNA-hSyn-GFP-KASH or AAV9-U6-scramble gRNA-hSyn-GFP-KASH were co-injected with AAV9-Mecp2-Cas9-HA. To determine whether *Npas4* regulates the memory generalization-discrimination balance, AAV9-hSyn-Cre-GFP or AAV9-hSyn-GFP were injected into *Npas4* conditional knockout mice.

### Stereotaxic viral injection

Mice were anesthetized with 1.5%–2% isoflurane in O_2_. Stereotaxic injections were performed unilaterally or bilaterally into the dorsal hippocampal DG with the following coordinates (relative to bregma) and volumes: AP -1.90 mm, ML ± 1.20 mm, DV -2.00 mm, 120-160 nL per hemisphere. Viruses were infused at a rate of 100 nL per minute and needles were kept at the injection site for 6 minutes. To target MEC and LEC, the following coordinates and volumes were used: MEC (AP -4.75 mm, ML ± 3.25 mm, DV -3.60 mm, 250-300 nL per hemisphere), LEC (AP -3.40 mm, ML ± 4.35 mm, DV -4.10 mm, 250-300 nL per hemisphere). For experiments with Dox-dependent ensemble labeling, animals were put on Dox diet (200 mg/kg, Bio-Serv) after surgery.

### Drug injections

To induce seizure, kainic acid (KA, Sigma) was dissolved in saline (1 mg/mL, Teleflex Medical) and injected i.p. at 20 mg/kg. Animals were sacrificed 1.5 hours later to check immediate early gene expression.

### Contextual fear conditioning (CFC)

Prior to CFC, mice were handled daily in a holding room for 3 days. For experiments with Dox-dependent ensemble labeling, the Dox diet was replaced by the regular diet (off Dox) after the last handling session on the third handling day. Behavioral assays were typically carried out 48 hours after the last handling session.

On the training day, mice were first transported to the holding room and allowed to acclimate for at least 30 minutes. Next, they were transported to the behavioral room and placed in context A: 24 cm (L) x 19 cm (W) x 17.5 cm (H), with steel grid floors and 1% acetic acid (Sigma). Mice were conditioned as follows: animals were allowed to explore the conditioning chamber freely for 4 minutes and received 2-s 0.5-mA foot shocks at 1-minute intervals 3 times, starting at the 58th second. After each experiment, the chamber was cleaned with 70% ethanol and subsequently with water. For experiments with Dox-dependent ensemble labeling, mice were switched back to the Dox diet 24 hours after CFC.

To evaluate contextual fear memory generalization and discrimination, mice were returned to the behavioral suite 48 hours post-conditioning for a series of 4-minute recall sessions. Context A’ was designed to be highly similar to the conditioning context A, retaining all physical features except for the substitution of the olfactory cue with 0.25% benzaldehyde (Sigma). Context B was designed to be moderately similar to the conditioning context A, differing by the addition of a blue foam pad floor insert and the removal of all olfactory cues. Context C was designed to be a highly distinct environment, using a different chamber with dimensions 32 cm (L) x 25 cm (W) x 32 cm (H) characterized by opaque white walls, a padded floor and a scent of 0.25% benzaldehyde (Sigma) under attenuated lighting conditions. The recall sessions were conducted over a two-day period with an inter-trial interval of at least 6 hours to minimize memory interference. On the first day, animals were exposed to either context A’ or context B, followed by re-exposure to context A. On the second day, animals were tested in the distinct context C, followed by a final re-exposure to context A. Contextual fear memory expression was measured by manually scoring freezing behavior over each 4-minute recall session, sampling every 5 s, with freezing defined as absence of movement for at least 1 s. Scorers were blind to experimental conditions. Freezing behavior was quantified across all sessions to determine whether mice generalized or discriminated between the conditioning and novel contexts.

### Immunohistochemistry and confocal imaging

Animals were drop fixed with 4% paraformaldehyde (PFA, Sigma) in phosphate buffered saline (PBS). Coronal sections were cut at 50 mm thickness. For immunohistochemistry experiments, brain slices were washed in PBS for 10 minutes 3 times and blocked for 2 hours at room temperature with the blocking solution: 1% Triton X-100 (Sigma) and 10% goat serum (Invitrogen) in PBS. The slices were then incubated with primary antibody overnight at 4 °C in the antibody solution: 0.1% Triton X-100 and 5% goat serum in PBS. The next day, the slices were washed in PBS for 10 minutes 3 times and incubated with secondary antibody for 2 hours at room temperature in the antibody solution. Finally, the slices were washed in PBS for 10 minutes 3 times and mounted with DAPI Fluoromount-G (Southern Biotech).

Antibodies used and dilutions were as follows: rabbit anti-Npas4 (1:10,000, home-made), rabbit anti-Fos (1:1,000, Santa Cruz), and Alexa 647 secondary antibodies (1:500, Invitrogen).

All images were acquired using a Zeiss LSM800 confocal microscope under a 10x objective. Analyses were performed by investigators who were blind to experimental conditions.

### Electrophysiology

Mice were subjected to electrophysiology 2-3 days after CFC. Mice were anesthetized and perfused with an oxygenated (95% O_2_, 5% CO_2_) ice-cold cutting solution containing (in mM): 213 sucrose, 1.25 NaH_2_PO_4_, 2 CaCl_2_, 3 KCl, 2 MgSO_4_, 26 NaHCO_3_, 11 D-Glucose, 1.3 sodium-L-ascorbate, and 0.6 sodium pyruvate (300 mOsm osmolarity, pH 7.30). After decapitation, the animal’s brain was rapidly dissected and transferred to the oxygenated ice-cold cutting solution. Coronal brain slices 300-µm thick were prepared with a Vibratome 1200S (Vibratome, USA) in the oxygenated ice-cold cutting solution. The slices were transferred to an incubation chamber with oxygenated artificial cerebrospinal fluid (ACSF), composed of (in mM) 125 NaCl, 1.25 NaH_2_PO_4_, 2 CaCl_2_, 3 KCl, 2 MgSO_4_, 26 NaHCO_3_, 11 D-Glucose, 1.3 sodium-L-ascorbate, and 0.6 sodium pyruvate (300 mOsm osmolarity, pH 7.30), for 30 min at 32 °C and then at room temperature (25 ± 2 °C) for > 30 min before use.

Whole-cell recordings from DG neurons in the acute slices were done with an Axonpatch 700B amplifier (Molecular Devices, USA), using an upright microscope (FN1, Nikon, Japan) equipped with infrared differential interference contrast (DIC) optics. Recording micropipettes were made from borosilicate glass capillaries (B-120-69-15, Sutter Instruments, USA) with a Sutter P97 puller. Micropipette resistances were 3–6 MΩ.

To record optically evoked excitatory postsynaptic currents (oEPSCs) or optically evoked inhibitory postsynaptic currents (oIPSCs) in the DG, 2-ms 470-nm LED stimulations at 0.5-1.0 mW/mm^2^, generated from a mounted LED (Thorlabs), were controlled by TTL input and delivered through the 40x objective. Paired whole-cell recordings were carried out on labeled and unlabeled neighboring neurons. oEPSCs were recorded in the presence of tetrodotoxin (TTX, 0.5 µM), picrotoxin (PTX, 50 µM),w and 4-aminopyridine (4-AP, 100 µM) from cells voltage clamped at -70 mV with a Cs-based internal solution containing (in mM): 130 CsMeSO_3_, 10 phosphocreatine, 1 MgCl_2_, 10 HEPES, 0.2 EGTA, 4 Mg-ATP, 0.5 Na-GFP, pH adjusted to 7.25 with CsOH, 295 mOsm osmolarity. oIPSCs were recorded in the presence of tetrodotoxin (TTX, 0.5 µM), DNQX (20 µM), APV (50 µM), and 4-aminopyridine (4-AP, 100 µM) from cells voltage clamped at -70 mV with a high-Cl internal solution containing (in mM): 103 CsCl, 12 CsMeSO_3_, 5 TEA-Cl, 10 HEPES, 4 Mg-ATP, 0.5 Na-GTP, 0.5 EGTA, 1 MgCl_2_, pH adjusted to 7.25 with CsOH, 295 mOsm osmolarity.

mEPSCs were isolated with tetrodotoxin (TTX, 0.5 µM) and picrotoxin (PTX, 50 µM), and recorded from cells voltage clamped at -70 mV using the Cs-based internal solution described above. mIPSCs were isolated with tetrodotoxin (TTX, 0.5 µM), DNQX (20 µM) and APV (50 µM), and recorded from cells voltage clamped at -70 mV using the high-Cl internal solution described above.

Data were analyzed using MiniAnalysis (Synaptosoft) and Clampfit 10.2 (Molecular Devices) by investigators who were blind to experimental conditions.

### Neuron model

We modeled leaky integrate-and-fire neurons with spike frequency adaption. The membrane voltage *U*_*i*_ of neuron *i* evolved according to:^33^

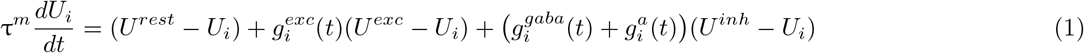

where τ^*m*^ denotes the membrane time constant, *U* ^*rest*^ denotes the membrane resting potential, *U* ^*exc*^ denotes the excitatory reversal potential, and *U* ^*inh*^ denotes the inhibitory reversal potential. The synaptic conductance terms 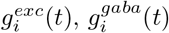, and 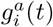 are described in the next section.

Neuron *i* fired a spike when its membrane voltage *U*_*i*_ surpassed the threshold *ϑ*_*i*_. Following a spike, *U*_*i*_ was set to *U* ^*rest*^ while *ϑ*_*i*_ was temporarily raised to *ϑ*^*spike*^. Without further spikes, *ϑ*_*i*_ decayed to its resting value *ϑ*^*rest*^ with time constant τ^*thr*^ following:^33^

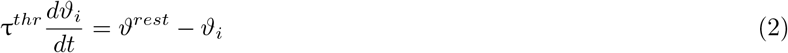

### Synapse model

We modeled conductance-based synapses. The inhibitory synaptic input 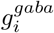 and the spike-triggered adaption *g*^*a*^ of neuron *i* evolved according to:^33^

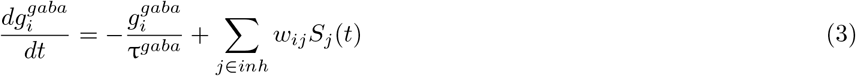

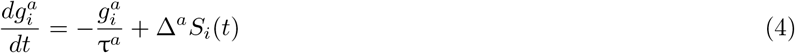

where 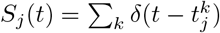 denotes the presynaptic spike train and 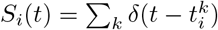 denotes the postsynaptic spike train. In each instance, *δ* denotes the Dirac delta function and 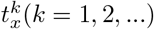 denotes the firing times of neuron *x. w*_*ij*_ denotes the weight of the synapse from neuron *j* to neuron *i*. Δ^*a*^ denotes the adaptation strength. τ^*gaba*^ denotes the GABA decay time constant and τ^*a*^ denotes the adaptation time constant.

The excitatory synaptic input of neuron *i* was determined by a combination of a fast AMPA-like conductance 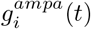 and a slow NMDA-like conductance 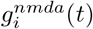:^33^

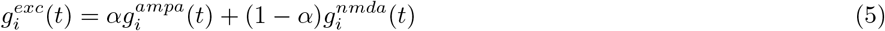

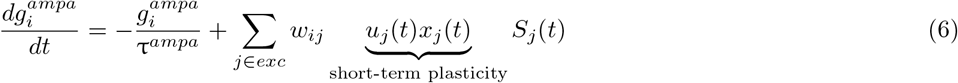

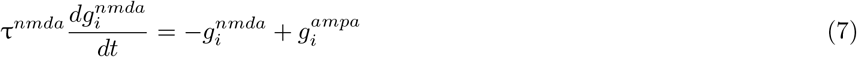

where *α* denotes a constant that sets the relative contribution of 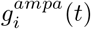 and 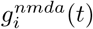 while τ^*ampa*^ and τ^*nmda*^ denote their respective time constants. *u*_*j*_(*t*) and *x*_*j*_(*t*) denote variables that reflect the state of short-term plasticity as described in the next section.

### Synaptic plasticity model

Our synaptic plasticity model built on previous work that combined excitatory and inhibitory forms of plasticity to enable stable memory encoding and recall in recurrent spiking neural networks.^33^

#### Short-term plasticity

The variables *u*_*j*_(*t*) and *x*_*j*_(*t*) reflecting the state of short-term plasticity evolved according to:^33^

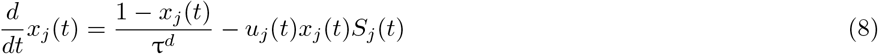

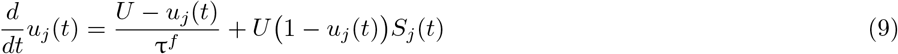

where τ^*d*^ and τ^*f*^ denote the depression and facilitation time constants, respectively. The constant *U* denotes the initial release probability.

#### Long-term plasticity

Upon initialization, the network model was subjected to a burn-in period to stabilize neural activity. During this period, we assumed that the entire population of excitatory neurons engaged both excitatory and inhibitory forms of long-term synaptic plasticity. This is consistent with the role of excitatory and inhibitory synaptic plasticity in the stabilization of neural activity during development.^29^ After the burn-in period, we assumed that half of the population of excitatory neurons expressed *Fos*-dependent excitatory synaptic plasticity (*Fos*^+^ neurons), while the other half expressed *Npas4* -dependent inhibitory synaptic plasticity (*Npas4*^+^ neurons). This is in line with (1) our previous study showing that *Fos*- and *Npas4* -dependent ensembles in mouse DG are distinct neuronal subpopulations,^17^ and (2) our experiments showing that these ensembles engage distinct forms of synaptic plasticity (Figure 1E/J).

We modeled excitatory synaptic plasticity during the burn-in period as a combination of triplet STDP,^30^ heterosynaptic plasticity,^31^ and transmitter-induced plasticity.^32^ Specifically, an excitatory synaptic weight *w*_*ij*_ from an excitatory neuron *j* to another excitatory neuron *i* followed:^33^

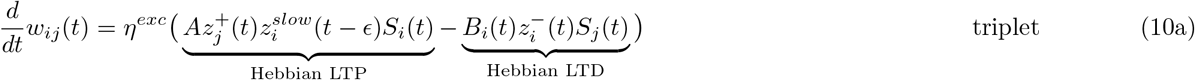

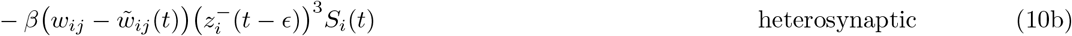

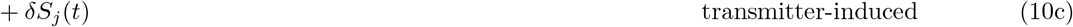

where *η*^*exc*^ denotes the excitatory learning rate, *A* denotes the LTP rate, *β* denotes the heterosynaptic plasticity strength, and *δ* denotes the transmitter-induced plasticity strength. *ϵ* denotes an infinitesimal offset whose purpose was to ensure that the current action potential was disregarded in the trace. State variables 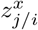 denote either pre- or postsynaptic traces with independent temporal dynamics set by a time constant τ^*x*^:^33^

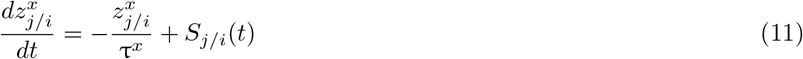

The reference weights 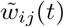 exhibited independent synaptic consolidation dynamics following the negative gradient of a double-well potential:^33^

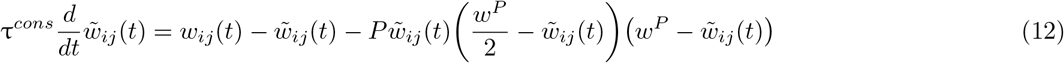

where *P* and *w*^*P*^ denote constants and τ^*cons*^ denotes the synaptic consolidation time constant. *P* set the magnitude of the double-well potential. When *w*^*P*^ = 0.5, an upper stable fixed point is located at 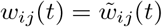 while a lower stable fixed point is located at 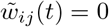. If 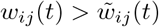 by a small margin, then the upper and lower stable fixed points of 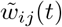 are increased. If 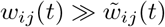, then 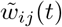 only retains a single fixed point that has a high value. This synaptic consolidation model is consistent with previous computational work^54^ and with synaptic tagging experiments that showed that long-lasting LTP is mediated by events taking place both during and prior to its initial induction.^55^ Furthermore, this model assumes that there are molecular mechanisms that enable synapses to maintain a stable efficacy (i.e., weight) even in the presence of intermittent fluctuations (e.g., triggered by molecular turnover).^56, 57^

The LTD rate *B*_*i*_(*t*) was subjected to homeostatic regulation:^33^

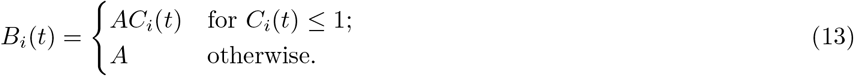

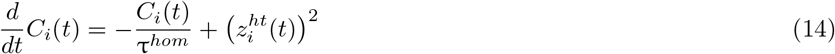

where τ^*hom*^ denotes a time constant and 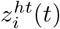 denotes a synaptic trace that follows Equation 11 with its own time constant τ^*ht*^.

We modeled inhibitory synaptic plasticity during the burn-in period as a form of homeostatic STDP. Specifically, an inhibitory synaptic weight *w*_*ij*_ from an inhibitory neuron *j* to an excitatory neuron *i* evolved according to:^33^

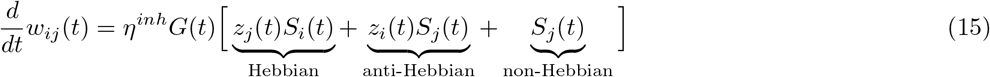

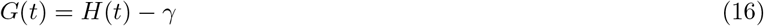

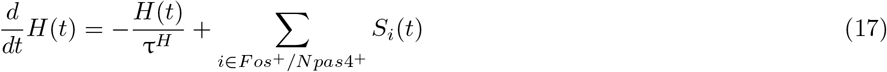

where *η*^*inh*^ denotes the inhibitory learning rate and *z*_*j/i*_ denotes either the pre- or postsynaptic trace following Equation 11 with a shared time constant τ^*iST DP*^ . *G*(*t*) tracks the difference between a hypothetical global secreted factor *H*(*t*) and the target activity level *γ* . *H*(*t*) denotes a low-pass-filtered version of the spikes fired by a population of excitatory neurons in the network with time constant τ^*H*^ . In particular, *H*(*t*) was computed separately for the population of neurons set to become *Fos*^+^ and the one set to become *Npas4*^+^ to ensure that both populations fired at the target rate at the end of the burn-in period. The fate of the postsynaptic neuron *i* after the burn-in period (i.e., *Fos*^+^ or *Npas4*^+^) determined which population of neurons was used to compute *H*(*t*). Note that Equation 15 is a unidirectional “depression-only” or “potentiation-only” inhibitory STDP rule: when the population activity level is below (*G*(*t*) *<* 0) or above (*G*(*t*) > 0) the target *γ*, the rule only allows depression or potentiation of inhibitory synapses, respectively. Therefore, the primary goal of inhibitory synaptic plasticity was to stabilize neural activity in a manner similar to previous theoretical models.^33, 34, 48^

After the burn-in period, we modeled *Fos*-dependent excitatory synaptic plasticity as a combination of Hebbian LTP and heterosynaptic plasticity. Specifically, an excitatory synaptic weight *w*_*ij*_ from an excitatory neuron *j* to a *Fos*^+^ excitatory neuron *i* followed:

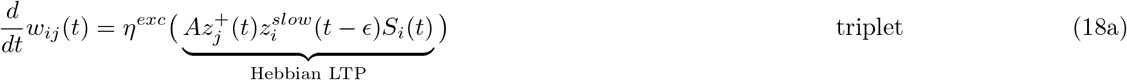

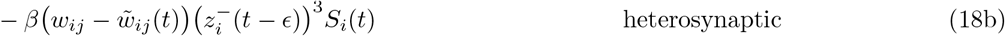

where we removed Hebbian LTD and transmitter-induced plasticity from Equation 10. 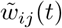 followed Equation 12.

After the burn-in period, we modeled *Npas4* -dependent inhibitory synaptic plasticity as an alternative form of homeostatic STDP. Specifically, an inhibitory synaptic weight *w*_*ij*_ from an inhibitory neuron *j* to an *Npas4*^+^ excitatory neuron *i* evolved according to:

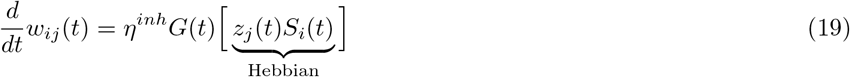

where we included only the Hebbian term and removed both the anti-Hebbian and non-Hebbian terms from Equation 15. *G*(*t*) and *H*(*t*) followed Equations 16 and 17, respectively.

In Figure S3 and Figure S4, we considered alternative formulations of either *Fos*-dependent excitatory synaptic plasticity (Equation 18) or *Npas4* -dependent inhibitory synaptic plasticity (Equation 19) and evaluated their effect on the computations promoted by *Fos*^+^ and *Npas4*^+^ ensembles. Where applicable, 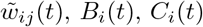, *G*(*t*), and *H*(*t*) followed Equations 12, 13, 14, 16, and 17, respectively.

Lastly, excitatory and inhibitory weights were restricted to a range defined by lower and upper bounds. Specifically, excitatory weights were bound between 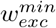 and 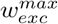, while inhibitory weights were bound between 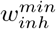 and 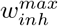 However, neither excitatory nor inhibitory weights reached their upper bound in our simulations, except in some cases where we considered alternative formulations of *Fos*- or *Npas4* -dependent synaptic plasticity.

### Network model

Our model consisted of an input population of *N*_*input*_ = 4, 096 Poisson neurons and a spiking neural network corresponding to DG. The DG network was composed of *N*_*exc*_ = 8, 192 excitatory neurons modeling granule cells and *N*_*inh*_ = 1, 024 inhibitory neurons modeling CCK^+^ interneurons. Feedforward excitatory synapses from the input population onto excitatory neurons had circular receptive fields centered at random locations (i.e., each excitatory neuron in the DG network received projections from a small circular area in the input population of radius *R*^*exc*^ whose random center location followed a uniform distribution). Feedforward synapses onto inhibitory neurons as well as recurrent synapses were randomly initialized following a uniform distribution with a probability of connection 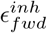 and *ϵ*_*rec*_, respectively. Feedforward synapses were initialized with specific weights (i.e., *w*^*input*→*E*^ and *w*^*input*→*I*^). Recurrent synapses were initialized with specific weights (i.e., *w*^*EI*^, *w*^*II*^, and *w*^*IE*^). For a complete list of model parameters, see Table S1.

During the burn-in period, synaptic plasticity was set as follows: (1) feedforward excitatory synapses onto excitatory neurons engaged short- and long-term excitatory synaptic plasticity, (2) feedforward and recurrent excitatory synapses onto inhibitory neurons engaged only short-term plasticity, (3) recurrent inhibitory synapses onto excitatory neurons engaged inhibitory synaptic plasticity, and (4) recurrent inhibitory synapses onto inhibitory neurons were static. After the burn-in period, synaptic plasticity changed in the following manner: (1) half of the population of excitatory neurons engaged *Fos*-dependent excitatory synaptic plasticity but no inhibitory synaptic plasticity (*Fos*^+^ neurons), (2) the other half of the population of excitatory neurons engaged *Npas4* -dependent inhibitory synaptic plasticity but no excitatory synaptic plasticity (*Npas4*^+^ neurons), (3) feedforward and recurrent excitatory synapses onto inhibitory neurons continued to engage only short-term plasticity, and (4) recurrent inhibitory synapses onto inhibitory neurons continued to be static. For a detailed description of synaptic plasticity in the model, see the previous section.

### Simulation protocols

Following our previous work,^35^ our simulation protocols consisted of multiple periods: burn-in, training, consolidation, probing, and recall. Each simulation began with a short burn-in period of duration *T*_*burn*_ that stabilized the activity of the DG network under background input from the input population at rate *ν*^*bg*^. Subsequently, the training stimulus was randomly presented to the network in the training period of duration *T*_*training*_. Next, the training stimulus was randomly reactivated in the consolidation period of duration *T*_*consolidation*_. This was motivated by previous experiments that showed that training-activated neurons in the entorhinal cortex of rodents and humans are selectively reactivated during post-training periods of sleep or rest.^36, 37^ At regular intervals over the course of the consolidation period (0, 1, …, *T*_*consolidation*_ h), the network advanced to a probing period of duration *T*_*probing*_ when the training stimulus was randomly presented (Figure 2B). This allowed us to identify the neuronal ensembles that encoded the training stimulus at the end of each consolidation interval (see the next section). Alternatively, the network proceeded to a recall period of duration *T*_*recall*_ at the end of each consolidation interval (0, 1, …, *T*_*consolidation*_ h) (Figure 3A). For each stimulus (i.e., either the training stimulus or one of three novel stimuli; Figure 3B), a separate recall period was carried out with partial cues of the given stimulus being presented. As a result, there were a total of four distinct recall periods (i.e., one for the training stimulus and one for each of the three novel stimuli) at the end of each consolidation interval. We depicted each stimulus and partial cue in a 64 x 64 grid (Figure 3B) and measured the intersection between the training stimulus and each novel stimulus (Figure 3C). When presenting or reactivating a stimulus or cue, off and on intervals were drawn from exponential distributions with means *T*_*off*_ and *T*_*on*_, respectively. During the presentation or reactivation of a stimulus or cue, the input neurons that corresponded to the given stimulus or cue selectively increased their firing rate to *ν*^*stim*^ while the remaining input neurons continued to fire at the background rate *ν*^*bg*^. For a complete list of simulation parameters, see Table S1.

### Neuronal ensembles in simulations

In our simulations, *Fos*^+^, *Npas4*^+^, and CCK^+^ neuronal ensembles were identified in the probing periods over the course of consolidation (see the previous section). Specifically, a neuron was considered part of the ensemble encoding the training stimulus at a given consolidation time point if its average stimulus-evoked firing rate during the ensuing probing period was above the activation threshold *ζ*^*thr*^ = 16 Hz. Note that a neuron may be part of the ensemble at an earlier time point but not at a later time point (i.e., the neuron drops out of the ensemble).^3, 35^ Conversely, a neuron may not be part of the ensemble at an earlier time point but part of it at a later time point (i.e., the neuron drops into the ensemble).^3, 35^ Furthermore, a neuron may alternate between being and not being part of the ensemble over the course of consolidation.^35^ Of course, a neuron may or may not be part of the ensemble at the end of the training period (i.e., 0 h of consolidation) and remain so throughout the consolidation period. Lastly, training stimulus neurons in the input population of our model were considered to be a stable ensemble (i.e., without turnover).

### Simulation and data analysis details

In our simulations, we employed the forward Euler method to update neuronal state variables with a time step Δ = 0.1 ms (with the exception of reference weights 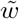 for which we set a longer time step Δ_*long*_ = 1.2 s for efficiency reasons). The average firing rate of a population of neurons in our model (i.e., *Fos*^+^, *Npas4*^+^, or CCK^+^) was computed with a temporal resolution of 10 ms without smoothing or convolution.

### Code details

Simulation code was written in C++ using the Auryn simulator for spiking neural networks.^58^ Following several preliminary simulations, we found that setting the number of Message Passing Interface (MPI) ranks *N*_*ranks*_ = 16 minimized our simulation time with Auryn. Code used to analyze simulation output was written in Python 3.10.

### Quantification and statistical analysis

#### Neuronal ensemble turnover

To examine the temporal evolution of *Fos*^+^, *Npas4*^+^, and CCK^+^ neuronal ensembles in our simulations, we computed the rate of overlap, drop-out, and drop-in between consecutive consolidation time points (Figure 2G-H and Figure S2C). Overlap was defined as the number of neurons part of both the ensemble at consolidation time t - 1 h and the ensemble at consolidation time t h divided by the total number of neurons in the union of the two ensembles. Drop-out was defined as the number of neurons part of the ensemble at consolidation time t - 1 h but not at consolidation time t h divided by the total number of neurons in the ensemble at consolidation time t - 1 h. Drop-in was defined as the number of neurons part of the ensemble at consolidation time t h but not at consolidation time t - 1 h divided by the total number of neurons in the ensemble at consolidation time t h.

#### Cue-evoked ensemble firing rate and activation

In our simulations, cue-evoked ensemble firing rate was defined as the average cue-evoked firing rate of individual neurons in the ensemble. An ensemble was considered activated upon presentation of a partial cue if its cue-evoked firing rate was above the activation threshold *ζ*^*thr*^ = 16 Hz. Cue-evoked ensemble activation was defined as the number of instances when the ensemble was activated by the presentation of a cue divided by the total number of cue presentations during the recall period. Cue-evoked ensemble firing rate and activation were computed separately for the training and novel stimuli.

#### Generalization-discrimination index (GDI) and engram computations

We defined a generalization-discrimination index (GDI) between two quantities q_1_ and q_2_ as:

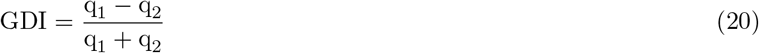

In our simulations, we computed a GDI between the ensemble activation evoked by recall cues of the training stimulus and a novel stimulus by setting q_1_ = activation_training stimulus_ and q_2_ = activation_novel stimulus_. activation_training stimulus_ and activation_novel stimulus_ denote ensemble activation when cues of the training stimulus or a novel stimulus were presented in the recall period of our simulation protocol, respectively. Note that a separate ensemble activation GDI was calculated between the training stimulus and each novel stimulus, with the resulting indices being averaged. We defined memory generalization and discrimination at the neural level as a change in ensemble activation GDI over the course of memory consolidation (i.e, between when ensembles emerged and 48 h of consolidation). Ensembles emerged at the end of training (i.e., 0 h of consolidation) in our simulations, except when we removed Hebbian LTP from *Fos*-dependent excitatory synaptic plasticity (Figure S3A-C). When activation GDI decreased over consolidation, the corresponding ensemble was said to support generalization. Conversely, when activation GDI increased over consolidation, the corresponding ensemble was said to support discrimination.

In our CFC experiments, we computed a GDI between freezing levels during recall sessions in the training context and a novel context by setting q_1_ = freezing_training context_ and q_2_ = freezing_novel context_. freezing_training context_ denotes freezing level in context A and freezing_novel context_ denotes freezing level either in context A’, B, or C. We defined memory generalization and discrimination at the behavioral level based on freezing behavior GDI. When freezing behavior GDI was not significantly different from zero, mice were said to generalize. Conversely, when freezing behavior GDI was significantly different from zero, mice were said to discriminate.

#### Statistical analysis

Statistical analyses were performed using Prism 10 (GraphPad Software). The means of two groups were compared using a Wilcoxon signed-rank test when the samples were matched. When there were two independent variables, the means of multiple groups were compared using a two-way analysis of variance (ANOVA) followed by Sidak’s multiple comparisons test. A one-sample *t* test was used to compare group means to hypothetical values. All comparisons were two-sided. Statistical significance was determined as follows: \**p <* 0.05, \*\**p <* 0.01, \*\*\**p <* 0.001. Unless otherwise noted, data were shown as mean ± standard error of the mean (SEM).

## Supplemental figures

**Figure S1.**
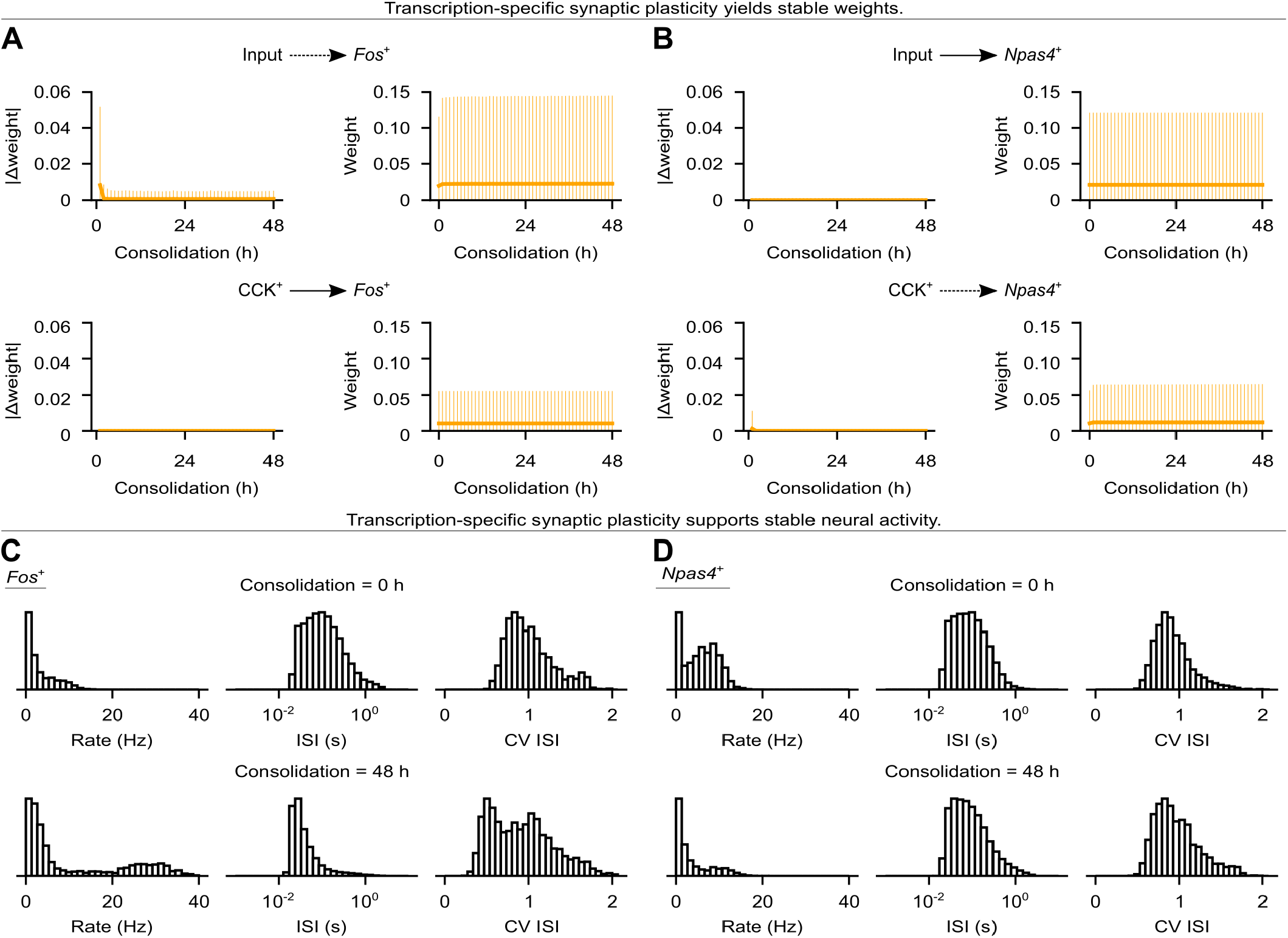
Analysis of synaptic weight strength and neural activity. (A-B) Post-encoding evolution of synapses onto *Fos*^+^ (A) or *Npas4*^+^ (B) neurons in Figure 2A when the network was subjected to the simulation protocol in Figure 2B. Top: feedforward excitatory synapses. Bottom: recurrent inhibitory synapses. Left: change in synaptic weight strength. Right: synaptic weight strength. Plasticity of synapses as in Figure 2A. Representative trial shown. Data shown as mean ± standard deviation (SD). (C-D) Analysis of the activity of *Fos*^+^ (C) or *Npas4*^+^ (D) neurons during the intervals marked at the top of Figure 2E and Figure 2F, respectively. Top: at 0 h of consolidation. Bottom: at 48 h of consolidation. From left to right: histograms of firing rates, interspike intervals (ISI), and coefficient of variation (CV) of ISI. Representative trial shown.

**Figure S2.**
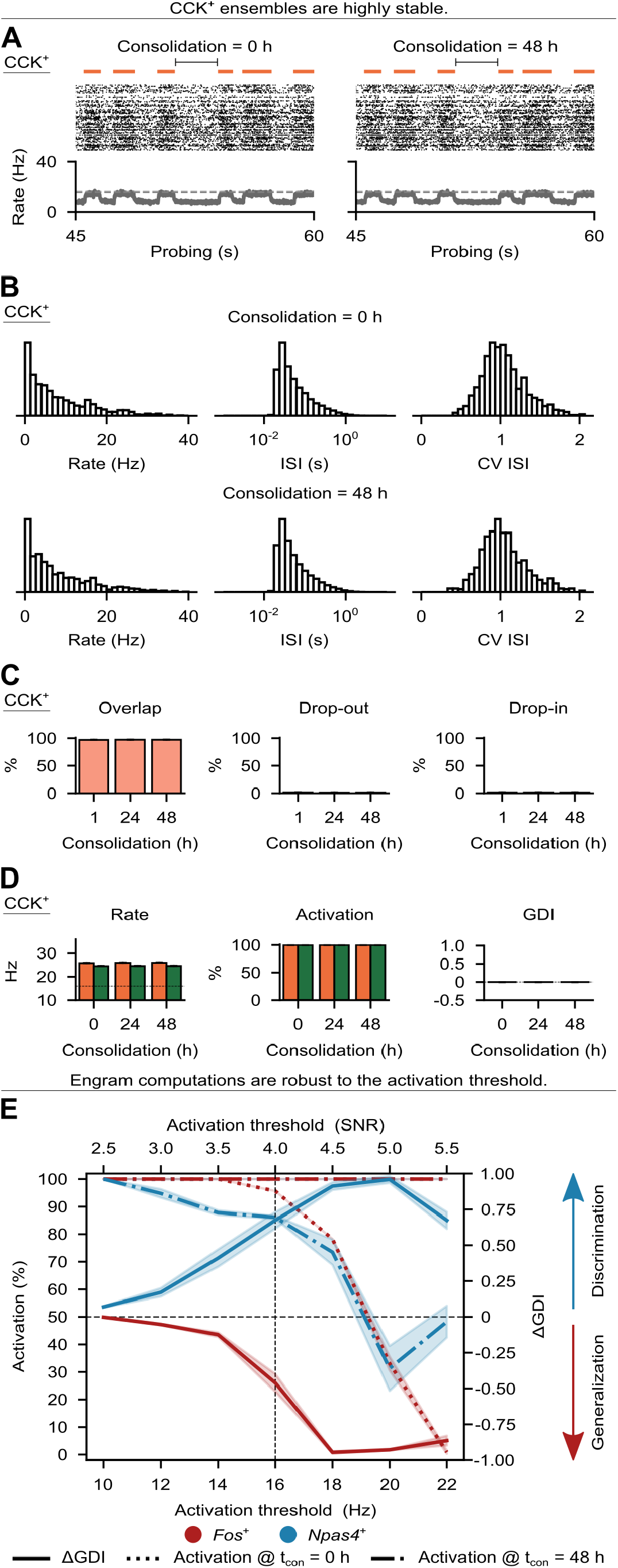
Analysis of CCK^+^ interneurons and sensitivity analysis of the activation threshold. (A-D) Analysis of CCK^+^ interneurons in Figure 2A when the network was subjected to the simulation protocols in Figure 2B and Figure 3A. (A) Activity of CCK^+^ interneurons during probing at 0 h (left) and 48 h (right) of consolidation. Top: training stimulus presentation times (in orange). Middle: spike raster of 256 randomly-chosen interneurons (for clarity, only every fifth spike is shown). Bottom: average firing rate of the population of interneurons (dashed line indicates the activation threshold *ζ*^*thr*^ = 16 Hz). Population rate shown without smoothing or convolution. Representative trial shown. (B) Analysis of the activity of CCK^+^ interneurons during the intervals marked at the top of (A). Top: at 0 h of consolidation. Bottom: at 48 h of consolidation. From left to right: histograms of firing rates, interspike intervals (ISI), and coefficient of variation (CV) of ISI. Representative trial shown. (C) Post-encoding evolution of CCK^+^ ensembles. Overlap (left), drop-out (middle), and drop-in (right) measured between consecutive time points as in Figure 2G-H (see Methods). (D) Analysis of CCK^+^ ensembles. Left: cue-evoked ensemble firing rate (dashed line indicates the activation threshold *ζ*^*thr*^ = 16 Hz). Middle: cue-evoked ensemble activation. Right: GDI between the ensemble activation evoked by cues of the training and novel stimuli. Color denotes stimulus as in Figure 3B. (E) Sensitivity analysis of ensemble activation and computations when the network in Figure 2A was subjected to the simulation protocols in Figure 2B and Figure 3A. Ensemble activation evoked by cues of the training stimulus at 0 and 48 h of consolidation and associated change in ensemble activation GDI as a function of the activation threshold. Threshold in units of signal-to-noise ratio (SNR) calculated as the firing rate threshold divided by the target firing rate for inhibitory synaptic plasticity *γ* = 4 Hz (see Methods). Vertical dashed line indicates the reference threshold used throughout the manuscript. Horizontal dashed line indicates activation = 50% and ΔGDI = 0. Data shown as mean ± SEM and *n* = 10 trials in (C), (D) and (E).

**Figure S3.**
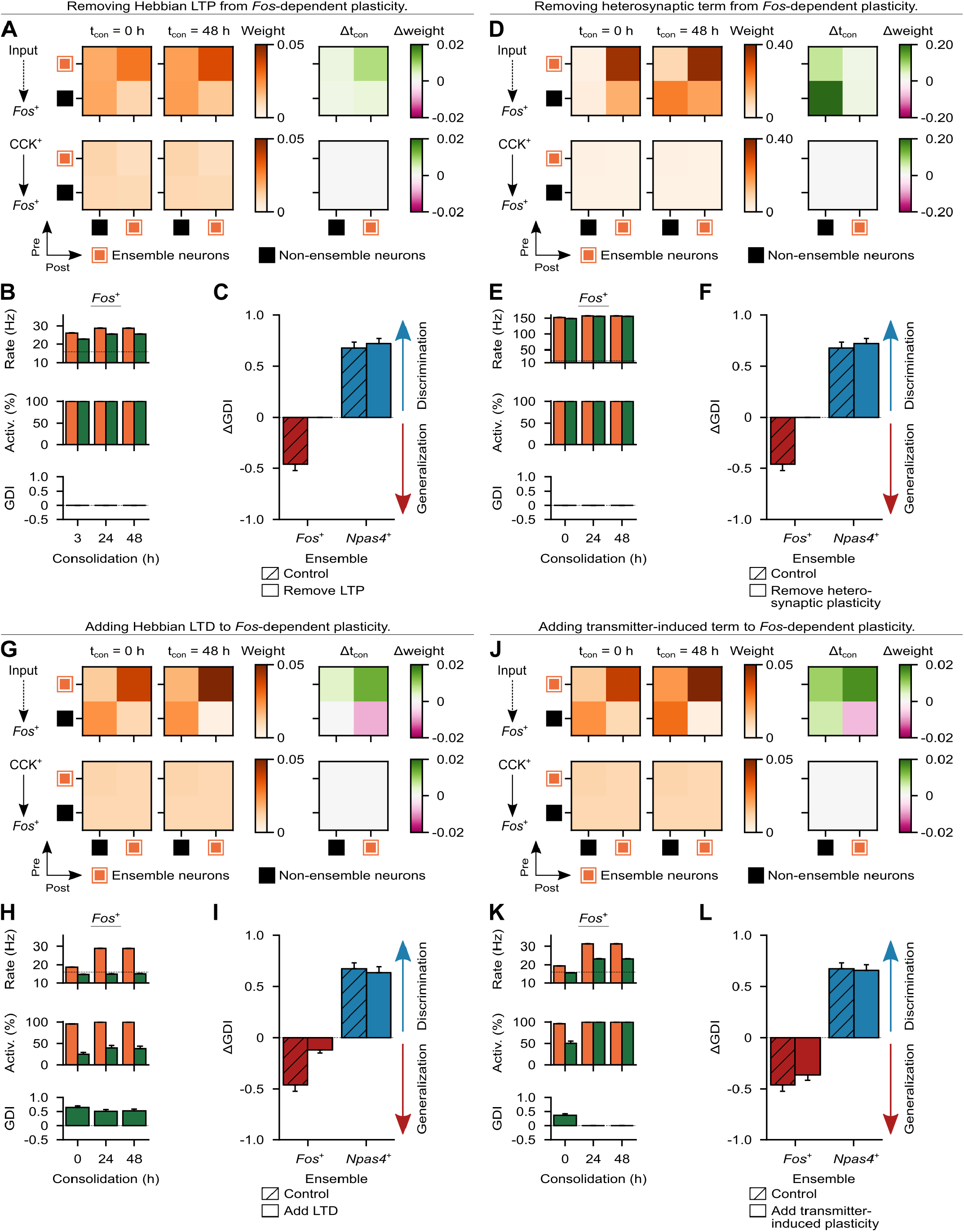
Analysis of alternative forms of *Fos*-dependent excitatory synaptic plasticity. (A-C) Analysis of the effect of removing Hebbian LTP from *Fos*-dependent excitatory synaptic plasticity when the network in Figure 2A was subjected to the simulation protocols in Figure 2B and Figure 3A (see Methods). (A) Mean weight strength of synapses onto *Fos*^+^ neurons clustered according to ensemble status (i.e., ensemble and non-ensemble neurons) at the end of training (i.e., 0 h of consolidation). Note that, when Hebbian LTP was removed from *Fos*-dependent plasticity, we identified *Fos*^+^ ensemble neurons at 0 h of consolidation considering the training period as opposed to the probing period following training. This was done because *Fos*^+^ ensemble neurons could only be identified considering probing periods starting at consolidation = 3 h. Top: feedforward excitatory synapses. Bottom: recurrent inhibitory synapses. Left: at 0 h of consolidation. Middle: at 48 h of consolidation. Right: change in weight strength between 0 and 48 h of consolidation. Plasticity of synapses as in Figure 2A. Representative trial shown. (B) Analysis of *Fos*^+^ ensembles. Top: cue-evoked ensemble firing rate (dashed line indicates the activation threshold *ζ*^*thr*^ = 16 Hz). Middle: cue-evoked ensemble activation. Bottom: GDI between the ensemble activation evoked by cues of the training and novel stimuli. Color denotes stimulus as in Figure 3B. Note that, when Hebbian LTP was removed from *Fos*-dependent plasticity, *Fos*^+^ ensembles emerged in probing periods after 3 h of consolidation, whereas *Npas4*^+^ ensembles emerged at 0 h of consolidation (i.e., at the end of training) (see Methods). (C) Change in ensemble activation GDI between when ensembles emerged in probing periods and 48 h of consolidation. A decrease in GDI indicates generalization, whereas an increase indicates discrimination. Control denotes the simulations with the default formulation of *Fos*-dependent plasticity (i.e., Hebbian LTP and heterosynaptic plasticity, see Methods) (same data as Figure 3E). (D-F) Same as (A-C) but for the analysis of the effect of removing the heterosynaptic term from *Fos*-dependent excitatory synaptic plasticity (see Methods). Note that, when the heterosynaptic term was removed from *Fos*-dependent plasticity, *Fos*^+^ and *Npas4*^+^ ensembles emerged at 0 h of consolidation (i.e., at the end of training) (see Methods). (G-I) Same as (A-C) but for the analysis of the effect of adding Hebbian LTD to *Fos*-dependent excitatory synaptic plasticity (see Methods). Note that, when Hebbian LTD was added to *Fos*-dependent plasticity, *Fos*^+^ and *Npas4*^+^ ensembles emerged at 0 h of consolidation (i.e., at the end of training) (see Methods). (J-L) Same as (A-C) but for the analysis of the effect of adding the transmitter-induced term to *Fos*-dependent excitatory synaptic plasticity (see Methods). Note that, when the transmitter-induced term was added to *Fos*-dependent plasticity, *Fos*^+^ and *Npas4*^+^ ensembles emerged at 0 h of consolidation (i.e., at the end of training) (see Methods). Data shown as mean ± SEM and *n* = 10 trials in (B-C), (E-F), (H-I), and (K-L).

**Figure S4.**
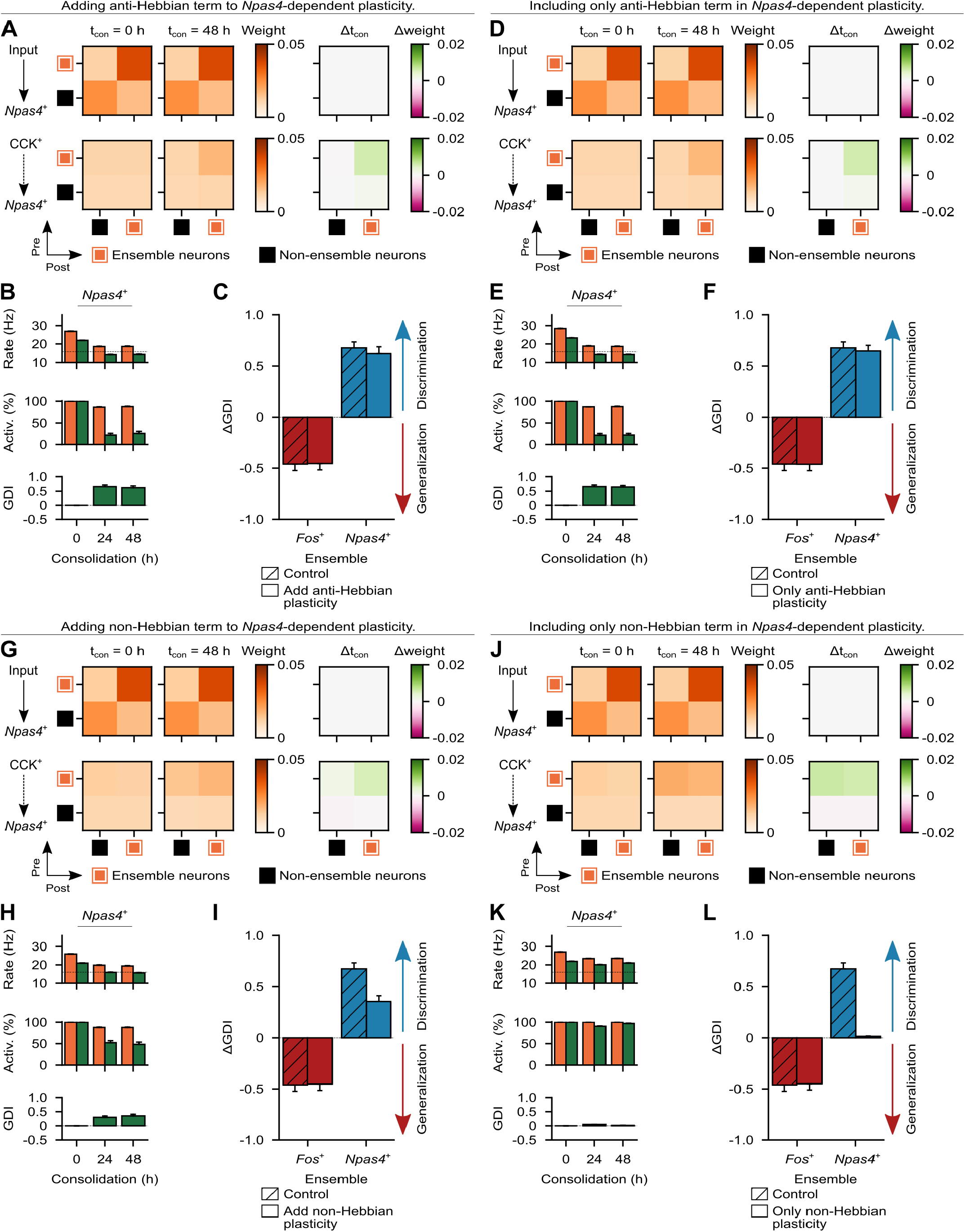
Analysis of alternative forms of *Npas4* -dependent inhibitory synaptic plasticity. (A-C) Analysis of the effect of adding the anti-Hebbian term to *Npas4* -dependent inhibitory synaptic plasticity when the network in Figure 2A was subjected to the simulation protocols in Figure 2B and Figure 3A (see Methods). (A) Mean weight strength of synapses onto *Npas4*^+^ neurons clustered according to ensemble status (i.e., ensemble and non-ensemble neurons) at the end of training (i.e., 0 h of consolidation). Top: feedforward excitatory synapses. Bottom: recurrent inhibitory synapses. Left: at 0 h of consolidation. Middle: at 48 h of consolidation. Right: change in weight strength between 0 and 48 h of consolidation. Plasticity of synapses as in Figure 2A. Representative trial shown. (B) Analysis of *Npas4*^+^ ensembles. Top: cue-evoked ensemble firing rate (dashed line indicates the activation threshold *ζ*^*thr*^ = 16 Hz). Middle: cue-evoked ensemble activation. Bottom: GDI between the ensemble activation evoked by cues of the training and novel stimuli. Color denotes stimulus as in Figure 3B. Note that, when the anti-Hebbian term was added to *Npas4* -dependent plasticity, *Fos*^+^ and *Npas4*^+^ ensembles emerged at 0 h of consolidation (i.e., at the end of training) (see Methods). (C) Change in ensemble activation GDI between when ensembles emerged and 48 h of consolidation. A decrease in GDI indicates generalization, whereas an increase indicates discrimination. Control denotes the simulations with the default formulation of *Npas4* -dependent plasticity (i.e., only the Hebbian term, see Methods) (same data as Figure 3E). (D-F) Same as (A-C) but for the analysis of the effect of including only the anti-Hebbian term in *Npas4* -dependent inhibitory synaptic plasticity (see Methods). Note that, when only the anti-Hebbian term was included in *Npas4* -dependent plasticity, *Fos*^+^ and *Npas4*^+^ ensembles emerged at 0 h of consolidation (i.e., at the end of training) (see Methods). (G-I) Same as (A-C) but for the analysis of the effect of adding the non-Hebbian term to *Npas4* -dependent inhibitory synaptic plasticity (see Methods). Note that, when the non-Hebbian term was added to *Npas4* -dependent plasticity, *Fos*^+^ and *Npas4*^+^ ensembles emerged at 0 h of consolidation (i.e., at the end of training) (see Methods). (J-L) Same as (A-C) but for the analysis of the effect of including only the non-Hebbian term in *Npas4* -dependent inhibitory synaptic plasticity (see Methods). Note that, when only the non-Hebbian term was included in *Npas4* -dependent plasticity, *Fos*^+^ and *Npas4*^+^ ensembles emerged at 0 h of consolidation (i.e., at the end of training) (see Methods). Data shown as mean ± SEM and *n* = 10 trials in (B-C), (E-F), (H-I), and (K-L).

## Supplemental table

**Table S1.** List of model and simulation parameters. Value source indicated as below: (a) values taken from Friedemann Zenke, Everton J Agnes, and Wulfram Gerstner. Diverse synaptic plasticity mechanisms orchestrated to form and retrieve memories in spiking neural networks. *Nature Communications*, 6(1):1–13, 2015. (b) values chosen somewhat arbitrarily without targeted optimization. (c) values optimized over several preliminary simulations.

| Neuronal populations |  |  |
| --- | --- | --- |
| Parameter | Value<sup>(source)</sup> | Description |
| $N_{exc}$ | 8192<sup>(b)</sup> | Size of the population of excitatory neurons in the DG network model. |
| $N_{inh}$ | 1024<sup>(a)</sup> | Size of the population of inhibitory neurons in the DG network model. |
| $N_{input}$ | 4096<sup>(a)</sup> | Size of the population of input neurons. |
| Network connectivity |  |  |
| Parameter | Value<sup>(source)</sup> | Description |
| $\epsilon_{fwd}^{inh}$ | 0.05<sup>(c)</sup> | Probability of connection of feedforward excitatory synapses onto inhibitory neurons. |
| $\epsilon_{rec}$ | 0.05<sup>(a)</sup> | Probability of connection of recurrent synapses. |
| $R^{exc}$ | 8<sup>(a)</sup> | Radius of the receptive field of excitatory neurons. |
| $w^{input \rightarrow E}$ | 0.5<sup>(a)</sup> | Initial weight of feedforward excitatory synapses onto excitatory neurons. |
| $w^{input \rightarrow I}$ | 0.4<sup>(c)</sup> | Initial weight of feedforward excitatory synapses onto inhibitory neurons. |
| $w^{EI}$ | 0.01<sup>(c)</sup> | Fixed weight of recurrent excitatory synapses onto inhibitory neurons. |
| $w^{II}$ | 0.2<sup>(a)</sup> | Fixed weight of recurrent inhibitory synapses onto inhibitory neurons. |
| $w^{IE}$ | 0.2<sup>(a)</sup> | Initial weight of recurrent inhibitory synapses onto excitatory neurons. |
| Neuron model |  |  |
| Parameter | Value<sup>(source)</sup> | Description |
| $\tau^m$ | 20 ms<sup>(a)</sup> | Membrane time constant. |
| $U^{rest}$ | -70 mV<sup>(a)</sup> | Membrane resting potential. |
| $U^{exc}$ | 0 mV<sup>(a)</sup> | Excitatory reversal potential. |
| $U^{inh}$ | -80 mV<sup>(a)</sup> | Inhibitory reversal potential. |
| $\tau^{thr}$ | 5 ms<sup>(a)</sup> | Threshold time constant. |
| $\vartheta^{rest}$ | -50 mV<sup>(a)</sup> | Threshold resting value. |
| $\vartheta^{spike}$ | 100 mV<sup>(a)</sup> | Threshold value immediately after a spike. |
| Synapse model |  |  |
| Parameter | Value<sup>(source)</sup> | Description |
| $\tau^{gaba}$ | 10 ms<sup>(a)</sup> | GABA decay time constant. |
| $\tau^a$ | 100 ms<sup>(a)</sup> | Adaptation time constant. |
| $\Delta^a$ | 0.1<sup>(a)</sup> | Adaptation strength. |
| $\alpha^E$ | 0.2<sup>(a)</sup> | AMPA/NMDA ratio for excitatory neurons. |
| $\alpha^I$ | 0.3<sup>(a)</sup> | AMPA/NMDA ratio for inhibitory neurons. |
| $\tau^{ampa}$ | 5 ms<sup>(a)</sup> | AMPA decay time constant. |
| $\tau^{nmda}$ | 100 ms<sup>(a)</sup> | NMDA decay time constant. |
| Short-term synaptic plasticity model |  |  |
| Parameter | Value<sup>(source)</sup> | Description |
| $\tau_{EE}^d$ | 150 ms<sup>(a)</sup> | Depression time constant for excitatory synapses onto excitatory neurons. |
| $\tau_{EI}^d$ | 200 ms<sup>(a)</sup> | Depression time constant for excitatory synapses onto inhibitory neurons. |
| $\tau^f$ | 600 ms<sup>(a)</sup> | Facilitation time constant for excitatory synapses. |
| $U$ | 0.2<sup>(a)</sup> | Initial release probability for excitatory synapses. |
| Excitatory synaptic plasticity model |  |  |
| Parameter | Value<sup>(source)</sup> | Description |
| $\eta^{exc}$ | $1 \times 10^{-3(a)}$ | Learning rate of excitatory synapses. |
| $\lambda^\beta$ | 50<sup>(a)</sup> | $\beta/\eta^{exc}$ ratio. |
| $\tau^{cons}$ | 20 min<sup>(a)</sup> | Synaptic consolidation time constant. |
| $\lambda^\delta$ | 0.02<sup>(a)</sup> | $\delta/\eta^{exc}$ ratio. |
| $A$ | 1<sup>(a)</sup> | LTP rate. |
| $\tau^+$ | 20 ms<sup>(a)</sup> | Time constant of presynaptic trace for excitatory synaptic plasticity. |
| $\tau^-$ | 20 ms<sup>(a)</sup> | Time constant of postsynaptic trace for excitatory synaptic plasticity. |
| $\tau^{slow}$ | 100 ms<sup>(a)</sup> | Time constant of slow postsynaptic trace for excitatory synaptic plasticity. |
| $P$ | 20<sup>(a)</sup> | Potential strength. |
| $w^P$ | 0.5<sup>(a)</sup> | Upper fixed point of reference weight potential. |
| $\tilde{w}$ | 0.0<sup>(a)</sup> | Initial reference weight. |
| $\tau^{hom}$ | 10 min<sup>(a)</sup> | Time constant of homeostatic regulation of LTD. |
| $\tau^{ht}$ | 100 ms<sup>(a)</sup> | Time constant of postsynaptic trace for homeostatic regulation of LTD. |
| $w_{exc}^{max}$ | 5.0<sup>(a)</sup> | Maximum excitatory synaptic weight. |
| $w_{exc}^{min}$ | 0.0<sup>(a)</sup> | Minimum excitatory synaptic weight. |
| **Inhibitory synaptic plasticity model** |  |  |
| **Parameter** | **Value<sup>(source)</sup>** | **Description** |
| $\lambda^\eta$ | 50<sup>(a)</sup> | $\eta^{exc}/\eta^{inh}$ ratio. |
| $\gamma$ | 4 Hz<sup>(a)</sup> | Target activity level for a population of excitatory neurons. |
| $\tau^{iSTDP}$ | 20 ms<sup>(a)</sup> | Time constant of pre- and postsynaptic traces for inhibitory synaptic plasticity. |
| $\tau^H$ | 10 s<sup>(a)</sup> | Time constant of global secreted factor. |
| $w_{inh}^{max}$ | 5.0<sup>(a)</sup> | Maximum inhibitory synaptic weight. |
| $w_{inh}^{min}$ | 0.0<sup>(a)</sup> | Minimum inhibitory synaptic weight. |
| **Input model** |  |  |
| **Parameter** | **Value<sup>(source)</sup>** | **Description** |
| $\nu^{bg}$ | 5 Hz<sup>(a)</sup> | Background firing rate of the input population. |
| $\nu^{stim}$ | 15 Hz<sup>(c)</sup> | Firing rate of input neurons matching a given stimulus or cue when it is presented. |
| $T_{on}$ | 1 s<sup>(a)</sup> | Mean stimulus-on interval. |
| $T_{off}$ | 2 s<sup>(a)</sup> | Mean stimulus-off interval. |
| **Network simulation** |  |  |
| **Parameter** | **Value<sup>(source)</sup>** | **Description** |
| $T_{burn}$ | 60 s<sup>(b)</sup> | Duration of the burn-in period. |
| $T_{training}$ | 60 s<sup>(c)</sup> | Duration of the training period. |
| $T_{consolidation}$ | 48 h<sup>(b)</sup> | Duration of the consolidation period. |
| $T_{probing}$ | 60 s<sup>(c)</sup> | Duration of the probing period. |
| $T_{recall}$ | 90 s<sup>(b)</sup> | Duration of the recall period. |
| $\Delta$ | 0.1 ms<sup>(a)</sup> | Time step for updating neuronal state variables except for reference weights $\tilde{w}$. |
| $\Delta_{long}$ | 1.2 s<sup>(a)</sup> | Time step for updating reference weights $\tilde{w}$. |
| $N_{ranks}$ | 16<sup>(c)</sup> | Number of MPI ranks in Auryn simulations. |

